# Transmission of mutated SARS-CoV-2 variants is favored by relatively prolonged infections due to delayed immunity

**DOI:** 10.64898/2026.09.15.751875

**Authors:** Katherine Owens, Pierce Radecki, Stefano Tempia, Anne von Gottberg, Cheryl Cohen, Eli Boritz, Joshua T. Schiffer, Daniel B. Reeves

**Affiliations:** Fred Hutchinson Cancer Center, Vaccine and Infectious Disease Division, Seattle, WA; University of Washington, Department of Applied Mathematics, Seattle, WA; Vaccine Research Center, National Institute of Allergy and Infectious Diseases, National Institutes of Health, Bethesda, MD; Partnership for International Vaccine Initiative, Task Force for Global Health, Atlanta, GA; Centre for Respiratory Diseases and Meningitis, National Institute for Communicable Diseases (NICD) of the National Health Laboratory Service (NHLS), Johannesburg, South Africa; School of Pathology, Faculty of Health Sciences, University of the Witwatersrand, Johannesburg, South Africa; School of Public Health, Faculty of Health Sciences, University of the Witwatersrand, Johannesburg, South Africa; University of Washington School of Medicine, Department of Medicine, Seattle, WA; University of Washington, Department of Global Health, Seattle, WA

**Author notes:** These authors contributed equally to this work (Joshua T. Schiffer, Daniel B. Reeves).

**Keywords:** within-host evolution, within-host phylodynamics, mathematical model, multi-scale model, transmission dynamics

## Abstract

SARS-CoV-2 evolution enhanced viral fitness and immune evasion, extending the COVID-19 pandemic and resulting in millions of excess deaths. Viral diversity is generated within infected individuals, yet the timing and interplay of viral and immunological forces that drive transmissible evolution are incompletely understood. We developed a multi-scale within host phylodynamic (WiPhy) model of SARS-CoV-2 infection which couples viral replication, innate and acquired immune responses, and viral mutation. We then validated the model against quantitative viral and phylodynamic metrics. Model output predicts that typical acute infections rapidly generate genetic diversity due to accumulation of minor variants which in most cases do not achieve sufficient concentrations for transmission. Delayed innate immune responses correlate with higher peak viral load and diversification, allowing higher transmission risk of the founder virus or with a novel variant that is equally or less fit. In contrast, the risk of transmitting a fitter variant is highest during the ~10% of infections in which viral loads remain sufficiently high for transmission after 10-14 days. In these cases, non-sustained innate and/or weak acquired immune responses allow sufficient time for selection of a variant with one or more fitness enhancing non-synonymous mutations. Across a simulated cohort of ~1500 individuals, 5% of transmission risk came from variants with enhanced fitness from nonsynonymous mutations, and 13% of simulated infections accounted for 90% of fitter variant transmission risk. Our results highlight how the timing and interplay of viral and immunological forces within a host create bottlenecks that severely limit between host evolution.

## INTRODUCTION

Most SARS-CoV-2 infections occur in immune competent individuals and are contained rapidly, with detectable viral loads lasting 1-3 weeks^1–4^. Studies of within-host SARS-CoV-2 evolution during acute infection demonstrate that viral diversification is common, but novel mutations usually remain at low levels and early transmission with a tight bottleneck further limits their spread to new hosts^5–9^. Only a minority of infected individuals generates a mutated variant that eventually predominates at a within-host level^10^, raising the question of which conditions predispose to this outcome.

To connect the myriad nonlinear and time varying forces driving within-host evolution to population-level evolution, specialized, multi-scale mathematical models trained on longitudinal viral load and sequence data are needed. Models must recapitulate serial data because cross-sectional snapshots do not capture how viral dynamics influence evolution^8,9,11,12^. Yet, most existing models do not comprehensively include all forces governing this evolution. While phylodynamic models effectively describe within-host evolution for multiple pathogens^13–16^ they assume fixed pathogen population size and ignore critical, expanding, multi-component immune responses. Viral-immune dynamic (VID) models decode non-linear, mechanistic relationships which are not apparent with statistical analysis alone and interpolate viral loads and immune measures between samples. Prior SARS-CoV-2 VID models inferred the timing and intensity of immune responses against the entirety of replicating viral populations and provided a platform for accurately simulating therapies^17–25^. Some models specified a small number of pre-existing viral subpopulations to study drug resistance^25^ or immune evasion^26^, or were developed as one module within a larger, epidemiologic multiscale simulation framework^27,28^. Yet, these tools did not characterize variant emergence and dynamics at realistic levels of diversity and during time-varying immune responses.

Here we update a framework previously used to study evolution during early HIV-1 infection^29^ to study the forces governing variant replacement during acute, self-resolving SARS-CoV-2 infections. In this setting, we now distinguish non-synonymous (NS) from synonymous (S) mutations to facilitate comparison to real longitudinal data, where many of the observed SARS-CoV-2 intra-host single nucleotide variants (iSNV) have been S mutations^30^.

Existing models for SARS-CoV-2 infection have also partially linked viral kinetic and evolution patterns within a person to population-level epidemiologic patterns. Genomic data from household studies was used to characterize SARS-CoV-2 transmission dynamics, by estimating variant-specific secondary attack rates^31^, generation times, and stringent transmission bottlenecks of one to eight viruses^30,32^. Mathematical models were developed to estimate transmission risk as a function of viral load in the upper respiratory tract and to link this parameter to superspreader events^23,33–36^. Output from a stochastic transmission-bottleneck model allowed probability estimates for transmission of *de novo* SARS-CoV-2 mutations as a function of bottleneck size and a variant-specific selection coefficient ^37^. Yet, none of these models linked immune responses to within host viral evolution and onward to between host transmission and evolution.

To fill this gap, we developed a mechanistic mathematical model of within-host SARS-CoV-2 evolution during acute, self-limited infection (<40 days). We calibrated the model to paired longitudinal viral load and sequencing samples from 18 infections in immunocompetent individuals. We found that a highly skewed fitness-effect distribution of NS mutations was necessary to recapitulate granular viral load and diversity trends. We then projected viral evolution trajectories of 1452 real infections for which total viral load was obtained from nasal specimens. This allowed us to correlate model-predicted viral kinetic quantities and immune response parameters with key evolutionary metrics. Model output defined characteristics of highly fit new variants with single base-pair mutations and predicted the impact of host immune responses on their emergence. Finally, we linked within-host evolution and transmission risk models to identify conditions required for population spread of novel variants. Our results suggest that acute, self-limited infections in immunocompetent individuals generate multiple diverse new variants. Yet, only a subset of “relatively prolonged” infections (14-40 days) due to delayed immune responses has sufficient duration for a fitter new variant to predominate at viral loads needed for transmission.

## RESULTS

### Within-host phylodynamic model for acute SARS-CoV-2 infection

We previously developed and trained an ordinary differential equation (ODE) model for the dynamics of total viral load in the upper respiratory tract during acute SARS-CoV-2 infection in humans^17^. The model tracks susceptible cells, infected cells in an eclipse and productive state, virions, cells that become refractory to infection based on innate immune signaling, and an adaptive immune response that enhances clearance of infected cells later in infection (**Fig 1A**). See **Methods** for equations, a full list of parameter definitions, and population parameter values for the ordinary differential equations model.^17^

**Fig. 1.**
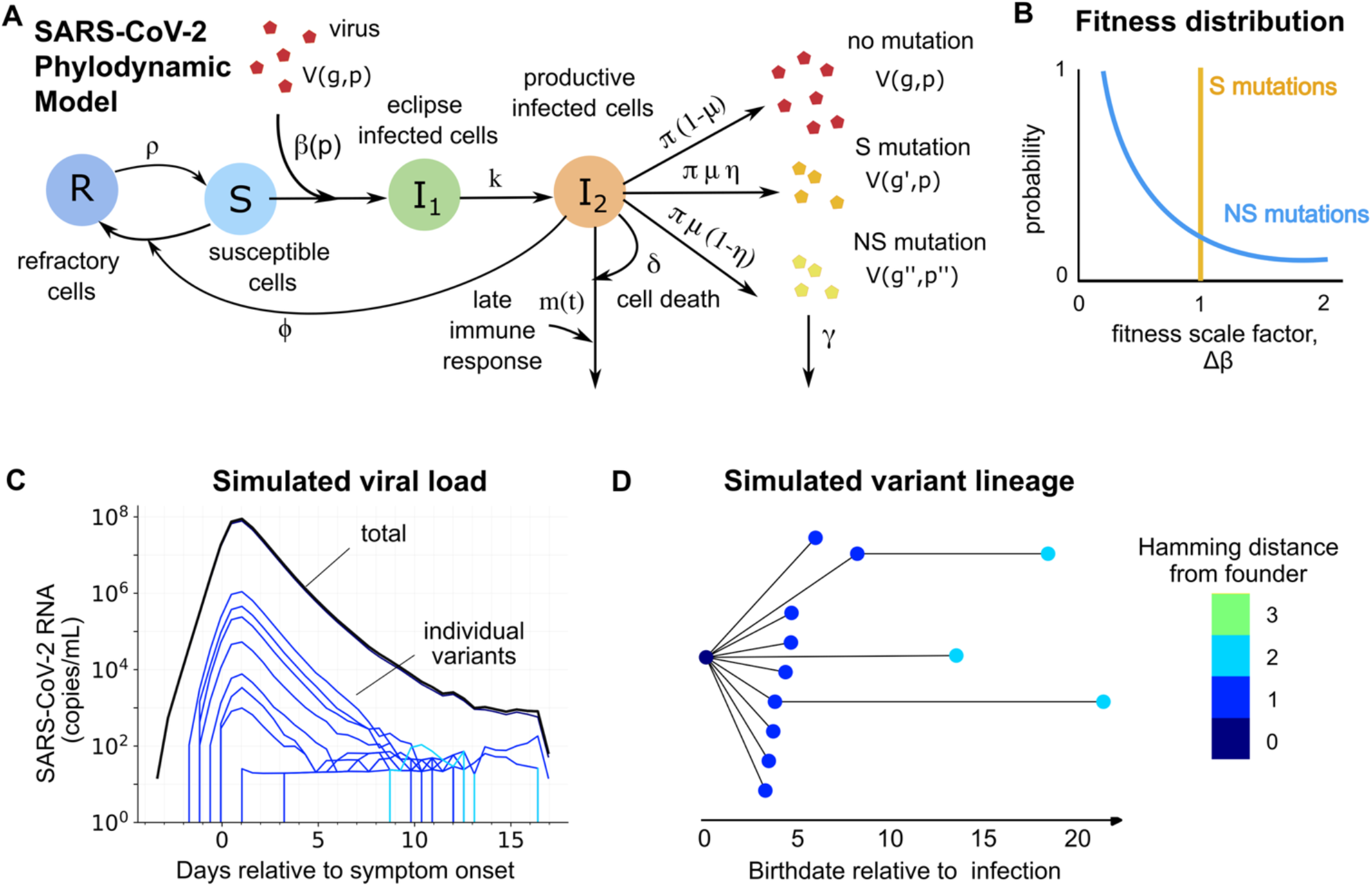
Within-host phylodynamic (WiPhy) model of SARS-CoV-2 infection. A) Wiphy mechanistic model. Susceptible cells S naturally proliferate and are infected by free viruses V(g, p), which each have a genotype (g, integer that specifies parental sequence and Hamming distance (HD) that specifies the number of basepair changes relative to founder virus) and phenotype (p, fitness quantified via infectivity parameter β(p)). Viruses are cleared naturally at rate γ. Infection proceeds through an eclipse phase I_1_ and into a productive phase I_2_ at which point new viruses can be produced at rate π. Susceptible cells can also enter a refractory state R based on infection levels (rate ϕ), and refractory cells can reenter the susceptible population (rate ρ). Productively infected cells die due to infection (rate δ) and also are cleared by an immune response m(t) that arises at time τ. New infection generates novel genotype variants at probability μ which are further distinguished by the gain of synonymous (no change in fitness, probability η) or non-synonymous mutations (fitness changed based on fitness model, probability 1 − η). B) WiPhy fitness model. Given a non-synonymous mutation, the fitness of the novel genotype variant g′ is the product of a fitness effect from an exponential distribution multiplied by the parental fitness β(p^′^) = Δβ × β(p). C) Simulated viral load. Model output tracks individual genotype variant viral loads over time (colored lines) that sum up to total viral load (black line). D) Simulated observed family tree. Downsampled model output can be used to construct a family tree relating variants (nodes) and ancestors (lineage connected with a line). In **C** and **D** viral loads and nodes on the family tree are colored by HD.

We updated the model into a within-host phylodynamic (WiPhy) framework to describe the evolutionary dynamics of self-limited SARS-CoV-2 infection. First, we considered the continuous-time Markov process for which the ODE model describes the behavior of the system in its mean field limit. Next, we model random mutations and fitness changes during viral replication via a branching process (**Fig 1A-B**). The stochastic model produces viral quasispecies, with each variant possessing a set of attributes: a unique identifier, a defining number of mutations, an infectivity, a birth date, and a parent variant from which it emerged. This allows simulation of an infection in which a variety of variants are created (**Fig 1C**). The transmission record of each variant is documented which allows us to generate a complete family tree relating all variants (**Fig 1D**). Model equations and the definitions of evolutionary parameters are in the **Methods**.

### Precise estimates of viral mutation rate and fitness distribution to recapitulate observed viral loads and variant evolution

We trained this WiPhy model on data from 18 individuals with well-documented acute infections who participated in one of two cohort studies of people diagnosed with COVID-19: a household cohort enrolled between October 2, 2020 and September 30, 2021^38^ or a hospitalized cohort enrolled between May 1, 2020 and December 31, 2020^39^. Data included a median of 4 (minimum 3, maximum 13) longitudinal measurements of viral load from upper respiratory tract swabs as well as longitudinal deep sequencing at a median of 3 (minimum 1, maximum 8) timepoints per individual. Sampling began a median of 4 (minimum −3, maximum 17) days post symptom onset and occurred every second day or three times weekly until viral clearance. Samples underwent high-throughput, single-genome amplification and sequencing (HT-SGS) of the full-length spike gene, an approach that combines unique barcoding of virus genomes with long-read sequencing to produce up to ~10^3^ single-copy sequences per sample^11^. See **Methods** for more details on the data used to calibrate the model. Throughout the text, we will refer to these 18 infected individuals as the South African Acute Infection (SAI) cohort.

We defined sixteen metrics that summarize both the viral load dynamics (peak, time to peak, time near peak and infection duration) and evolutionary dynamics (number of variants at days 3, 5, 10 and 20; average pairwise distance at days 3, 5, 10 and 20; frequency of the first dominant variant at days 3, 5, 10 and 20) of the virus over the course of infection as model-fitting targets (**Table S1, Fig S1A** and **Methods)**.

We assumed that the mutation rate and distribution of advantageous mutations is shared across individuals, though viral dynamic parameters vary. We fixed the fraction of S mutations at *f*_*s*_ = 0.35 based on previous estimates in the literature^11,40,41^, used a previously estimated distribution of viral dynamic parameters, and performed a grid search across three parameters: mutation rate (*μ*), average fitness change of NS variants (Δ*β*), and average hamming distance between a NS variant and its parent (Δ*hd*). For each candidate tuple of evolutionary parameters, we drew 30 sets of viral dynamic parameters from our previous modeling of the National Basketball Association (NBA) infection cohort (n = 1452)^17^ and ran 30 corresponding WiPhy simulations. Then, we randomly down sampled the WiPhy simulation outputs in time and sequence depth to accurately match the resolution of the data and calculated the 16 metrics defined in **Fig S1A** for the down-sampled model output (**Methods**). We assessed how closely model-generated output resembled SAI cohort data by applying the Kolmogorov-Smirnov test for each metric. Each metric that returned a non-significant difference between model and data garnered a point, for a maximum score of matching 16/16 metrics (**Fig S1B**). The process of down-sampling and scoring the model output against data was repeated 10 times to identify the parameter values with the best average score. We identified 6 parameter sets that on average matched at least 15/16 and one parameter set that matched all 16 metrics (**Fig S1C**).

The optimal match to the data was generated with a mutation probability (*μ*) of 0.02 per infection event with NS mutation events accompanied by an average of 1.5 distinguishing mutations (⟨*Δhd*⟩) and an average NS fitness multiplier (⟨*Δβ*⟩) of 0.2 given the exponential fitness distribution shape in **Fig 1B**. There was agreement (assess using two-sided Kolmogorov-Smirnov test) between the longitudinal data and optimal model output for viral load, number of variants detected, average pairwise distance, and frequency of the first dominant variant **(Fig 2A)** as well as comparisons of the distribution of the 16 metrics (**Fig 2B**).

**Fig. 2.**
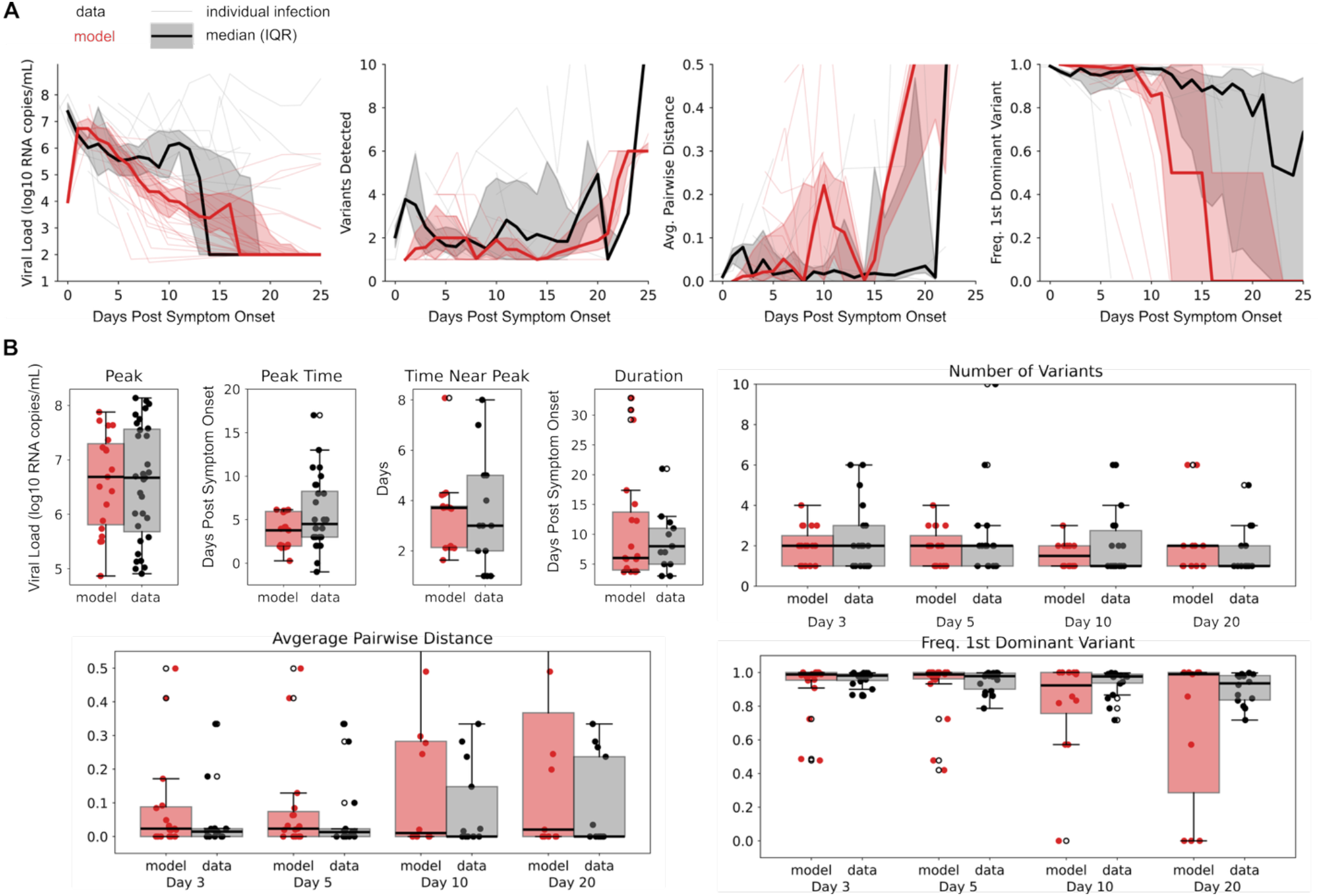
WiPhy model calibrated to viral load and phylodynamic data in SAI cohort. A) Agreement between model ouput with optimal parameter estimates (red lines) compared to the South African Acute Infection (SAI) cohort data (gray lines), The bold line indicates the median value (bold) and the shaded region indicates the 25^th^ and 75^th^ percentile (shading). From left to right, panels show the upper respiratory tract viral load, number of variants detected, average pairwise distance between sampled sequences, and frequency of the first dominant variant. B) Model output (red) was not significantly different from the SAI data from X individuals (black) with respect to 16 pre-defined metrics: peak viral load, time to peak viral load, time near (within 1 log) of peak viral load, infection duration, average pairwise distance at day 3, 5, 10 and 20 post infection, number of variants detected at day 3, 5, 10 and 20 post infection, and the frequency of the first dominant variant at day 3, 5, 10 and 20 post infection. Agreement was assessed by a two-sided Kolmogorov-Smirnov test (p<0.05).

### Simulated synonymous mutations required to generate realistic numbers of detectable minor variants

We next attempted model fitting while assuming exclusively NS mutations with exponentially distributed fitness effects. In this case, the model output either underestimated the number of variants detected or overestimated average pairwise distances relative to the data and consistently underestimated the persistence of the first detected variant. The maximum score attained by the model without S mutations was only an average of 14/16 metrics. In contrast, the model including S mutations generated many variants with identical fitness, which increased the chance of variant co-existence, a critical phenomenon that increases the number of detectable variants during periods in which the founder variant is dominant (**Fig 2A, right panel**).

### Agreement of model predicted viral fitness distributions with in vitro deep mutational scanning data

We next compared the model predicted fitness distribution that achieved best fit to the SAI cohort data with the distribution predicted by experimental deep mutational scanning (DMS) of the SARS-CoV-2 spike protein^42^. We assumed that 35% of mutations were S with fitness identical to the parental variant, while the remaining mutations were NS with fitness effect drawn from an exponential distribution with average ⟨Δβ⟩=0.2. The predicted distribution aligned closely with the DMS data^42^ (**Fig S2A**). The fraction of S mutations with identical fitness resulted in a horizontal line at 0 on the rank-log2 fitness effect curve from the calibrated evolutionary model. This simplifying assumption deviated somewhat from the experimental rank-log2 fitness effect curve, but the experimental curve intersected 0 within this region (**Fig S2A**). The cumulative distribution function for fitness showed that the fraction of mutations predicted to be non-deleterious by the model was 35.5% versus 21.3% in the DMS data (**Fig S2B**). This discrepancy between model and data arose because the DMS data includes a higher proportion of slightly deleterious and slightly beneficial mutations, than our fitness distribution permits. As we show later, slightly beneficial mutations rarely generate selective sweeps making this discrepancy acceptable. The two curves also reflect that most mutations are highly deleterious, agreeing that ~65% of mutations decrease fitness by 10% or more. Moreover, both the model and data project that highly beneficial mutations are exceedingly uncommon. In the empirical distribution only 0.3% of mutations confer a fitness advantage of 1.5 times the baseline fitness, and the model distribution predicts even fewer at 0.02%.

### Model generated realistic individual viral and evolutionary trajectories

After determining the optimal evolutionary parameters to match the SAI cohort, we used a population nonlinear mixed effects approach to match total viral loads and estimate individual viral dynamic parameters for the 18 individuals in the SAI cohort (**Methods**). We then ran 10 stochastic WiPhy simulations with population estimates for evolutionary parameters and the 18 individualized sets of viral dynamic parameters. For each of the 10 replicates, we also randomly sampled sequences from the simulation output at each time point 10 times (3 examples in **Fig 3**, all 18 in **Fig S3**). Because we did not vary the 3 parameters governing fitness and mutational distributions across individuals, our accurate fits to evolutionary metrics were emergent model properties achieved only by matching viral kinetics. We noted a higher degree of variability across different WiPhy simulations (distinct colors in **Fig 3**) than across different instances of sampling the same simulation (each line in **Fig 3**). This suggests that infection with very similar viral load profiles can generate different mutational profiles. Overall, these results allow us to project realistic evolutionary dynamics from infections with documented serial viral loads but no sequencing.

**Fig. 3.**
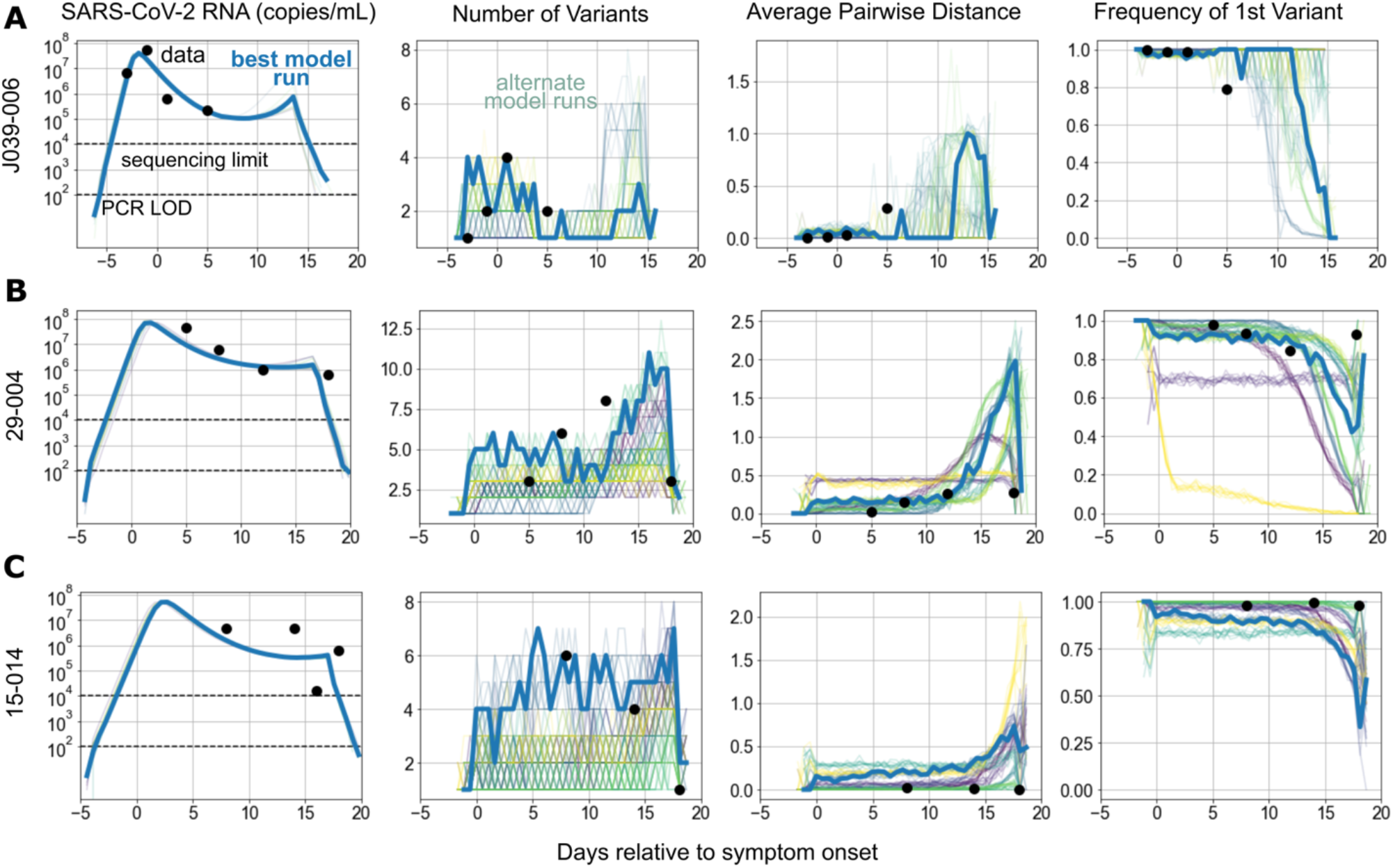
Individual phylodynamic output for participants generated using population-calibrated model. A-C) Comparison of model (lines) and data (dots) from three participants in the SAI cohort. From right to left: viral load, number of detectable variants, average pairwise genetic diversity of detected variants, and the frequency of the first dominant variant. Viral dynamic model parameters in each example were chosen by best fit to viral load data. 10 replicate simulations (colors) with 10 each stochastic subsamples (thin lines of same color) with bold blue line highlighting the sampled trajectory that best matched the four metrics.

### Correlation of peak viral load and viral load area under the curve (AUC) with within-host diversity

To determine relationships connecting viral dynamics and evolution, we performed a sensitivity analysis using infection dynamics from the NBA cohort. We used viral dynamic parameters from 1452 individual infections to simulate WiPhy trajectories and correlated 5 resulting viral load metrics to 8 resulting evolutionary metrics (**Fig 4A**). We found strong associations between peak and AUC viral load (log10 scale) and sequence diversity (maximum number of variants generated) and divergence (maximum hamming distance (HD) reached by a novel variant). Outlier infections with high divergence were evident at AUC>10^7^ RNA cp/mL x day **(Fig 4B)**. These results indicate that high viral peak drives diversification by increasing the absolute number of minor variants. Prolonged infection duration (>3 weeks) further increases the chance of sequential mutations occurring.

**Fig. 4.**
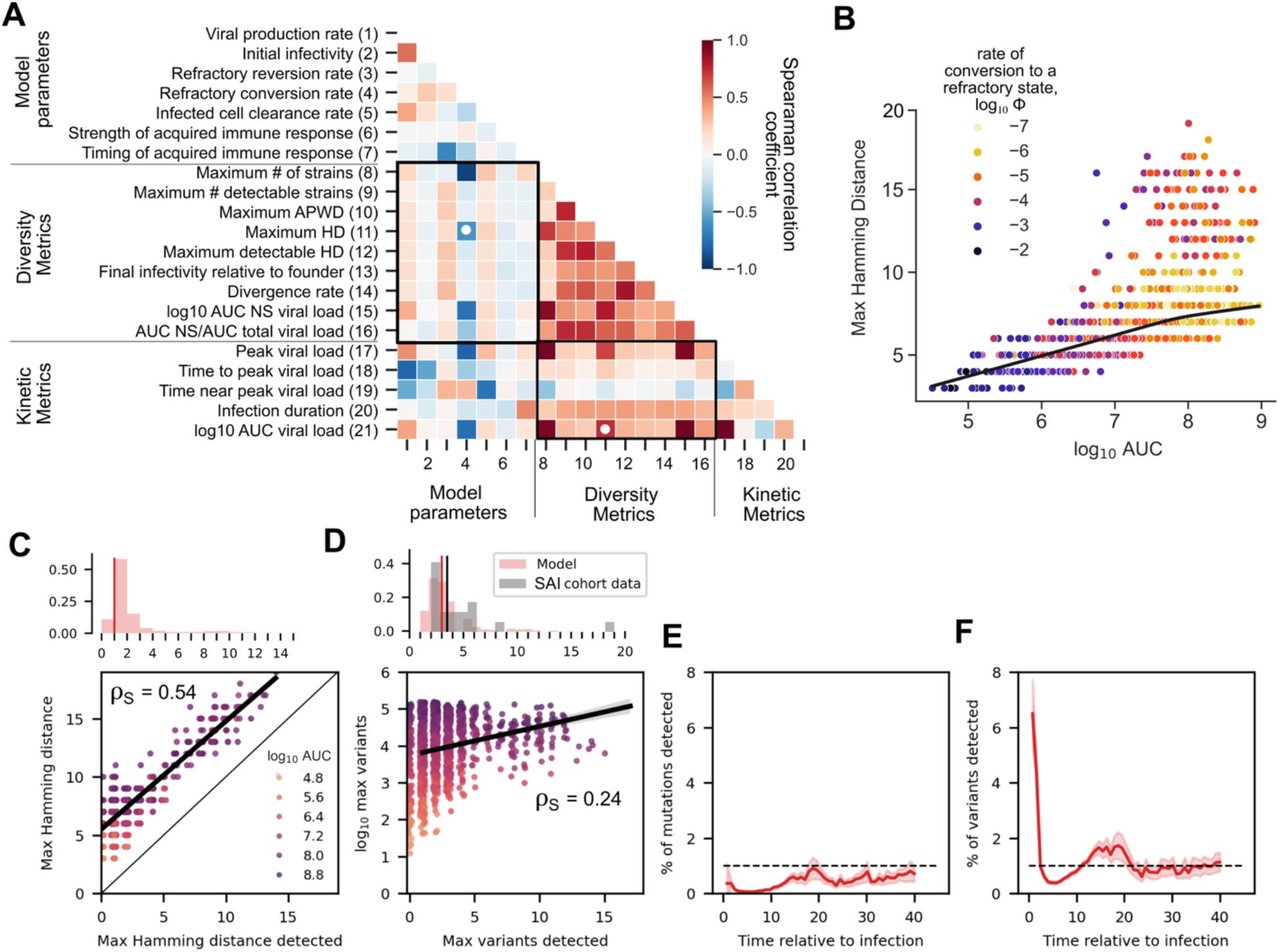
Correlation between model parameters, viral dynamic quantities, and diversity metrics. A) Sensitivity Analysis. The association matrices (Spearman correlation coefficient) between all input individual model parameters, model output diversity metrics, and model output viral load kinetic metrics calculated from simulations with best-fit viral dynamic parameters estimated from 1452 participants from NBA cohort and population calibrated evolutionary parameters (**Methods**). White dots indicates key tripartite association between refractory cell conversion rate, maximum Hamming Distance (max HD), and log10 area under the curve (AUC) viral load. B) Key strong positive relationship between log10 AUC viral load and max HD (metric 11 and 21, marked in panel A). Each dot represents simulation output based on one NBA cohort participant and black line is local non-parametric regression model across participants (LOWESS curve). Dots are colored by the log10 refractory conversion rate (ϕ). Values of ϕ range from a slower/weaker (−7, yellow) to a rapid/stronger (−2, navy) innate response. C) Strong association between max HD in the full simulation vs detected max HD in sampled simulation. Each dot represents simulation output based on one NBA cohort participant and bold line indicates linear relationship with high correspondence but lower concordance. Spearman correlation is noted. Upper inset histogram of detected HD values illustrates >50% of samples have max detected HD<5. D) Weak association between number of variants in full simulation vs detected number of variants in sampled simulation. The number of variants in the full simulation ranges from 10 to 100,000, scaling with the number of infected cells, whereas the detected number of variants in the sampled simulation ranges from 0 to 20. Each dot represents simulation output based on one NBA cohort participant and bold line indicates linear relationship with Spearman correlation noted. Upper inset histogram of detected variants illustrates >50% of samples have <5 variants in simulation (red bars), which matches with experimental data from the SAI cohort (gray bars). E&F) Percent of mutations and variants detected over time across 1452 simulations from the NBA cohort. Line is mean and shaded area is 95% CI.

### Delayed innate responses as a predicter of more rapid and extensive viral evolution

We next sought to determine which viral and immune parameters most strongly influenced viral evolution. We found the maximum HD, maximum number of variants, and maximum area under the curve of nonsynonymous variants were strongly negatively correlated with the rate at which susceptible cells become protected from infection in the presence of infected cells (*ϕ*) (**Fig 4A-B**). This model parameter was intended to indirectly capture the extent and efficiency of interferon signaling as an innate defense. A larger rate pushes susceptible cells to a refractory state earlier. This parameter was also strongly negatively associated with viral peak and AUC (**Fig 4A**). Other weaker correlations between viral dynamic parameters and evolutionary quantities were identified as well (**Fig 4A**). Viral production rate and the rate of reversion from a refractory state to susceptible state positively correlated with all measures of diversity and evolution. Overall, the refractory cell conversion rate was the parameter with strong correlation to viral burden and divergence (**Fig 4B**).

### Biased estimates of viral divergence and diversity in human studies due to lack of sequencing depth

To determine whether the true extent of SARS-CoV-2 evolution in an infected person is detected in a typical experimental sample, we assessed each simulated participant in the NBA cohort assuming experimental (frequency as low as 0.45%) versus complete (100% of variants) sequencing depth. We found a strong correlation between true and detected maximum HD (Spearman correlation *ρ*_*S*_ = 0.54, **Fig 4C**) but low concordance (Concordance Correlation Coefficient CCC = 0.22) as sampling consistently led to underestimation of true maximal HD by ~5 mutations. Most observed HD values were in the range of 0-1 mutations indicating limited evolution among dominant variants during most infections. Among simulations that reached HD >1 with experimental sampling, the correlation and concordance were stronger with *ρ*_*S*_ = 0.79 and CCC = 0.39.

There was only a weak association between number of variants detected with experimental versus complete sequencing depth (assessed on a linear scale). The number of variants was so under sampled that we were required to plot the true number of variants generated on a logarithmic scale against the number of variants detected on a linear scale (**Fig 4D**). We noted strong agreement between down-sampled model output and experimental distributions of detected variants in the SAI cohort data. Among infections with detected HD ≥1 and number of variants ≥2, AUC was often >10^6.5^ RNA cp/mL x day, corroborating that higher viral load drives diversification and increases the chance that a minor variant will be detected.

Calibrated^40,41^ model runs suggest that most *de novo* mutations and variants remain undetected throughout the course of infection. Across simulations, we found that at any given time less than 1% of model predicted mutations (**Fig 4E**) and less than 7% of unique variants (**Fig 4F**) were detected. The percent of detected variants fell to ~1% after peak viremia (**Fig 4F**).

### Stepwise increases in viral fitness during SARS-CoV-2 infection

Each WiPhy simulation generated detailed longitudinal data on all variants, including viral load, fitness, and ancestry. The dynamics of viral diversity in three infections are shown in **Fig 5** and **Fig S4-5**. We selected a simulation that demonstrated persistent infection (duration > 40 days) in **Fig 5** to visualize extensive within host evolution, relative to more typical rapidly contained infections in which variant generation was more limited (**Fig S4-5**). Based on a mutation rate of 0.02 new variants per cellular infection event and the average number of infection events exceeding 2.5 × 10^6^ cells per infection, each simulation generated tens of thousands of new variants. We assumed that only variants exceeding a frequency of 2% or greater were detectable provided that viral load was sufficiently high for sequencing. In each panel **Fig 5** and **Fig S4-5**, the founder variant is dark blue while variants with increasing HD (number of mutations) from the founder are shown in warmer colors.

**Fig. 5:**
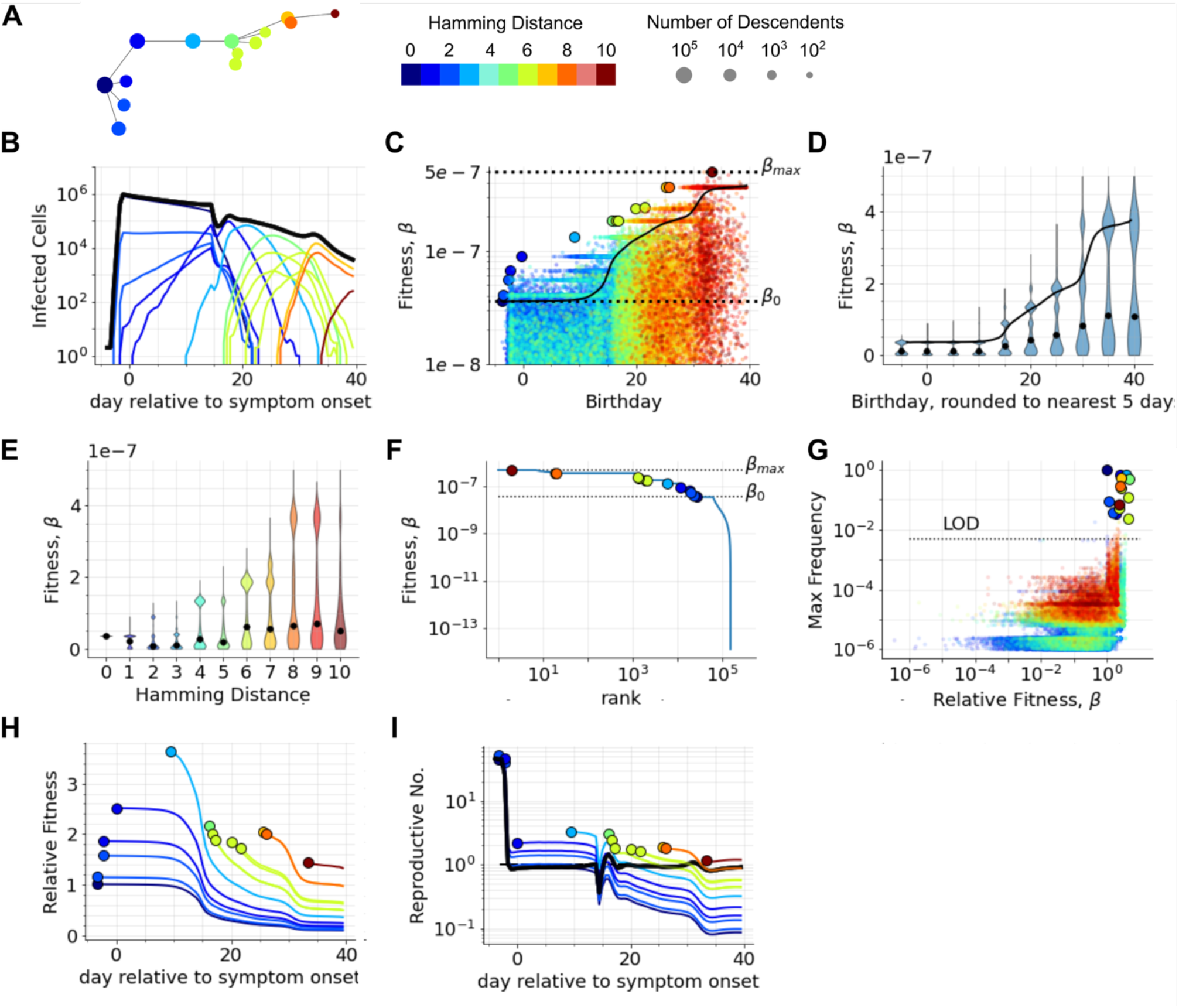
A detailed look at fitness evolution during infection. Panels A-I were generated using model output from one WiPhy simulation using population-calibrated evolutionary parameters and viral dynamic parameters estimated based on data from infection in an individual with unknown immune health. A) Simulated lineage of major variants (those reaching ≥2% of viral load during infection). Node radius is proportional to number of descendants (log10) and horizontal position corresponds to variant birthday (first time it appeared in simulation relative to symptom onset) concurrent with panel B. This color bar indicates HD for all panels **A-J**. B) Simulated number of productive infected cells (I_2_) corresponding to variants in **A** and their total (bold black). C) Relationship between variant birthday and absolute variant fitness. Single variants (small dots colored by HD) show heterogeneity and mean (black line) trails the fitness of emerging major variants (larger dots). Dashed horizontal lines denote initial variant fitness β_0_ and maximum β_max_ fitness allowed in the simulation. D) Violin plots of fitness of emerging variants grouped by 5 day intervals indicate bimodal fitness distributions with growing difference between mean (black line) and median (black dot). Note linear scale in comparison to **C**. E) The bimodal fitness distributions of variants grouped by HD show increasing variance as HD increases. Median value indicated by black dot. F) Each variant’s fitness plotted by rank (ordered from highest to lowest infectivity). Line shows complete distribution with large dots indicating major variants. Horizontal lines following major variants reffect synonymous variants. Many variants with fitness higher than the founder never reach detectable levels. G) Maximum frequency reached by variants versus their fitness relative to the population mean at time of variant appearance is roughly log log linear. Large dots indicate major variants above 2% frequency. These and other detectable minor variants (frequency above the limit of detection, LOD) have relative fitness near or above 1. H) Time dynamics of the relative fitness of major variants showing that relative fitness is highest when a strain emerges and then decreases – note linear scale in comparison to **E** in which many variants have small but heterogeneous fitness. I) Effective reproductive number (average number of newly infected cells from one infected cell, **Methods**) for each major variant (dot at birthday with line showing dynamics) compared to the population average (black line) over time after symptoms. New variants only can predominate when R>1.

For each WiPhy simulation, we constructed a family tree connecting each variant to its parent (**Fig 5A, Fig S4-5A**) that serves as a comparator to within-host phylogenies inferred from sequencing data. Unique to the WiPhy output, each node was sized according to the number of descendant variants that it generated. Tips are always smaller than their ancestral internal nodes, as their progeny is also part of the lineages of their ancestors. Eighteen days after symptom onset during the persistent infection simulation, the founder was eclipsed by a variant that had acquired one beneficial mutation shortly after peak viral load. Multi-generational progeny of this variant (light blue, turquoise, and gold) subsequently swept the population while the infection was still ongoing but trending towards clearance at day 40 when the simulation ended (**Fig 5B**). In more typical acute infections, novel variants emerged but never overtook the founding variant (**Fig S4B)** or overtook the founding variant near the end of infection (**Fig S5B**)

The fitness of detectable variants generated over time remained relatively constant (**Fig S4C**) or sequentially increased (**Fig 5C, S5C**). Newly detectable variants (larger circles) had substantially higher fitness than the average fitness of the viral population at the time of their birth or had a similar fitness but emerged very early in the viral expansion phase. There was a delay of 7-15 days between birth and detection as a fitter variant expanded and slowly increased mean variant fitness. Multiple horizontal bands of equally fit variants emerged from variants that reached 2% frequency, indicating blooms of S variant quasi-species (**Fig 5C, Fig S4-5C**). Grouping variants by birthdate rounded to the nearest 5 days revealed bimodal fitness distributions at each stage of infection (**Fig 5D, Fig S4-5D**). The majority of variants remained worse than the current average, with the gap between median and mean fitness growing as infection proceeded. We next compared the fitness distribution of variants conditioned on the genetic distance from the founder (**Fig 5E, Fig S4-5E**). The median variant fitness initially decreased, though the distribution was bimodal with an upper cluster of sequences near the current average fitness. In this example, the median fitness of variants remained lower than the founder fitness until the HD was 6. By day 15 when most of these more divergent variants were being generated, 3 beneficial mutations were fixed in the population.

### Sufficiently high relative fitness prerequisite for variant detection

In this example, only 13 of greater than 10000 high fitness variants achieved a detectable concentration (**Fig 5F**). Each high frequency variant had a cloud of associated variants with 1-2 synonymous variants at the same fitness rank that never reached a high frequency.

We next quantified variant characteristics required to reach a detectable frequency and found that the relative fitness at appearance (**Fig 5G, Fig S4-5G**), rather than the absolute fitness of a variant was more predictive of whether it would become detectable. All detectable variants had a relative fitness at the time of their birth of greater than or equal to 1, though this was not sufficient to guarantee reaching a detectable frequency or high frequency rank. Rather, a higher-than-average fitness at the time of appearance was a prerequisite to have a *chance* of reaching detectable levels with most highly fit new variants never exceeding 0.01% of total viruses (**Fig 5G**). This observation holds across infections.

The relative fitness of each variant peaked at the moment of its generation (**Fig 5H**). Soon after appearing, a fitter than average variant that increased in frequency augmented the average fitness of the viral population as well, thus decreasing its own relative fitness. As the variant came to dominate, it was responsible for a greater share of infection events, increasing the chance that one of its progeny would accrue a beneficial mutation and surpasses it in fitness, leading to a rapid decrease in its relative fitness to below one.

To exceed 2% frequency, each variant needed to appear during a stage of infection that allowed for an effective within-host reproductive number greater than 1 (**Fig 5I, Fig S4-5I**) which in turn depended not just on variant fitness, but also on the global immune pressure against all variants. Variants that appeared later during infection, when fewer target cells were available and acquired immune pressure was present, required a substantial fitness advantage over the founder variant to maintain a reproductive number only slightly greater than 1. In contrast, variants with a small fitness advantage that appeared early during infection had a much higher reproductive number than later variants due to the favorable conditions for viral expansion. Emerging variants also needed to avoid stochastic extinction and an infection duration long enough for the fitness advantage to manifest and reach detectable levels. Overall, variant selection was driven by rare variants with high relative fitness which emerged when immune pressure was present but not sufficient to eliminate all virions.

### Rarity and underestimation of novel variants takeover in human cohort studies

We defined novel variant “takeover” in infection as the moment when the frequency of the founder variant decreased below 50%. Assuming viral dynamic parameters from the NBA cohort, a takeover occurred in ~20% of simulated infections (**Fig S6A**). On average, variant takeover occurred 6 days after peak viral load, when viral load had decreased substantially. However, the distribution of takeover time relative to peak was wide, with a non-negligible minority of cases in which it occurred prior to peak viral load (**Fig S6B**).

Initially, this result seems to contradict results from the SAI cohort study which showed persistence of a single founder variant throughout SARS-CoV-2 infection in every individual without HIV^11^. This could simply follow from the relatively small sample size. Our simulations also show that pre-peak takeover could be misclassified as persistence of the founder variant: if the novel variant took over prior to the first sample, then the first *detected* variant would falsely appear to predominate throughout. Moreover, cases in which variant takeover occurred after viral load dropped below the minimum level required for sequencing were unlikely to be detected.

### Viral load AUC as a predictor of variant takeover

We next stratified the NBA cohort simulations based on six previously identified viral dynamic shedding patterns (**Fig S6C**) and compared the incidence of novel variant takeover in each group. Variant takeover was considerably less common in viral dynamic clusters 1 and 2 which had low viral AUC than in cluster 6 which had high viral AUC due to higher peak viral load and sustained replication (**Fig S6D**).

The timing of variant takeover somewhat correlated with later onset of the acquired immune response. In most cases with less than 20-day duration, novel variant taker over preceded model-predicted initiation of the acquired immune response. In these cases, the acquired response increased infected cell clearance and rapidly eliminated infection despite the increased within-host fitness of the novel variant (**Fig S6E**). In contrast, in specific circumstances when the acquired immune response initiated 5-10 days after symptoms and ~5 days before variant takeover, then infection had a higher chance of reaching an extended duration.

The extent to which viral infectivity increased over the course of infection varied widely, even among infections with novel variant takeover. The ratio of the final average viral infectivity to the initial infectivity was weakly correlated with infection duration. This association was largely driven by persistent infections (duration > 40 days) (**Fig S6F**). These persistent infections usually occurred in individuals with a fast rate of reversion from refractory back to susceptible state (darker purple dots in **Fig S6F**). In contrast, infections in which the protection afforded by the innate immune response was durable were unlikely to have extensive viral evolution that led to persistent infection. Together these observations suggest that a less durable innate immune response makes individuals vulnerable to accrual of fitness enhancing NS mutations, which, if followed by a weak acquired immune response, permits subsequent accrual of additional beneficial fitness mutations driving prolonged infections.

We further investigated the differences between simulations that remained founder dominant (founder frequency remains > 50%), cases of viral takeover that cleared (founder frequency <50% and viral load undetectable within 40 days) and cases of viral takeover that persisted (founder frequency <50% and viral load detectable through day 40). We compared viral dynamic metrics (**Fig S7A-E**), viral dynamic model parameter values (**Fig S7F-K**), and viral diversity metrics (**Fig S7L-O**) between these three groups. Even in cases that did not lead to viral persistence, the median infection duration was 3 days longer (p < 0.001, two-sided Mann-Whitney test) in cases of viral takeover than founder-dominant infections (**Fig S7A**). The median viral peak and log10 area under the viral load curve was also higher in the two takeover groups than in the founder-dominant group (**Fig S7B-C**), while the median time near the peak viral load was elevated in only the persistent takeover group (**S7D**). Time to peak viral load did not vary significantly among the three groups (p>0.01, two-sided Mann-Whitney test) (**S7E**).

Though the refractory cell conversion rate was strongly negatively correlated with viral diversity metrics, its median value was similar between these three groups (**S7F**). Takeover events occurred across moderate levels of refractory protection. However, as noted previously, the durability of the refractory state was a key differentiator between cases of takeover that were cleared versus those that persisted (**S7G**). Persistent takeover occurred in individuals with target cells rapidly cycling through the refractory state. Both takeover groups were associated with higher viral production rate compared to founder-dominant infections (**S7H**). Persistent takeover cases had a lower magnitude of the acquired immune response that initiated relatively early (**S7I-J**). In contrast, cases of viral takeover that cleared, had an elevated median death rate of infected cells (**S7K**).

As expected, viral diversity was higher and evolution of increased infectivity more pronounced in cases of variant takeover compared with founder dominated infections, and these metrics were higher still in persistent infections (**S7L-N**). The median area under the NS variant viral load curve was higher for cases of viral takeover, but many founder-dominated strains had comparably high area under the NS variant viral load curves (**S7O**).

### High proportion of novel variant transmission with synonymous mutations at peak viral load

We projected the risk of transmitting novel variants using a previously calibrated SARS-CoV-2 transmission risk model (**Fig 6A**). In the example shown, a novel variant took over around a week after the peak viral load (**Fig 6B**). If a contact event occurred, the absolute risk of transmitting a novel variant was ~10% at both peak viral load and 8 days post-peak (**Fig 6C**). Yet, the relative risk of transmitting a variant versus founder was much higher at day 8 post peak.

**Fig. 6:**
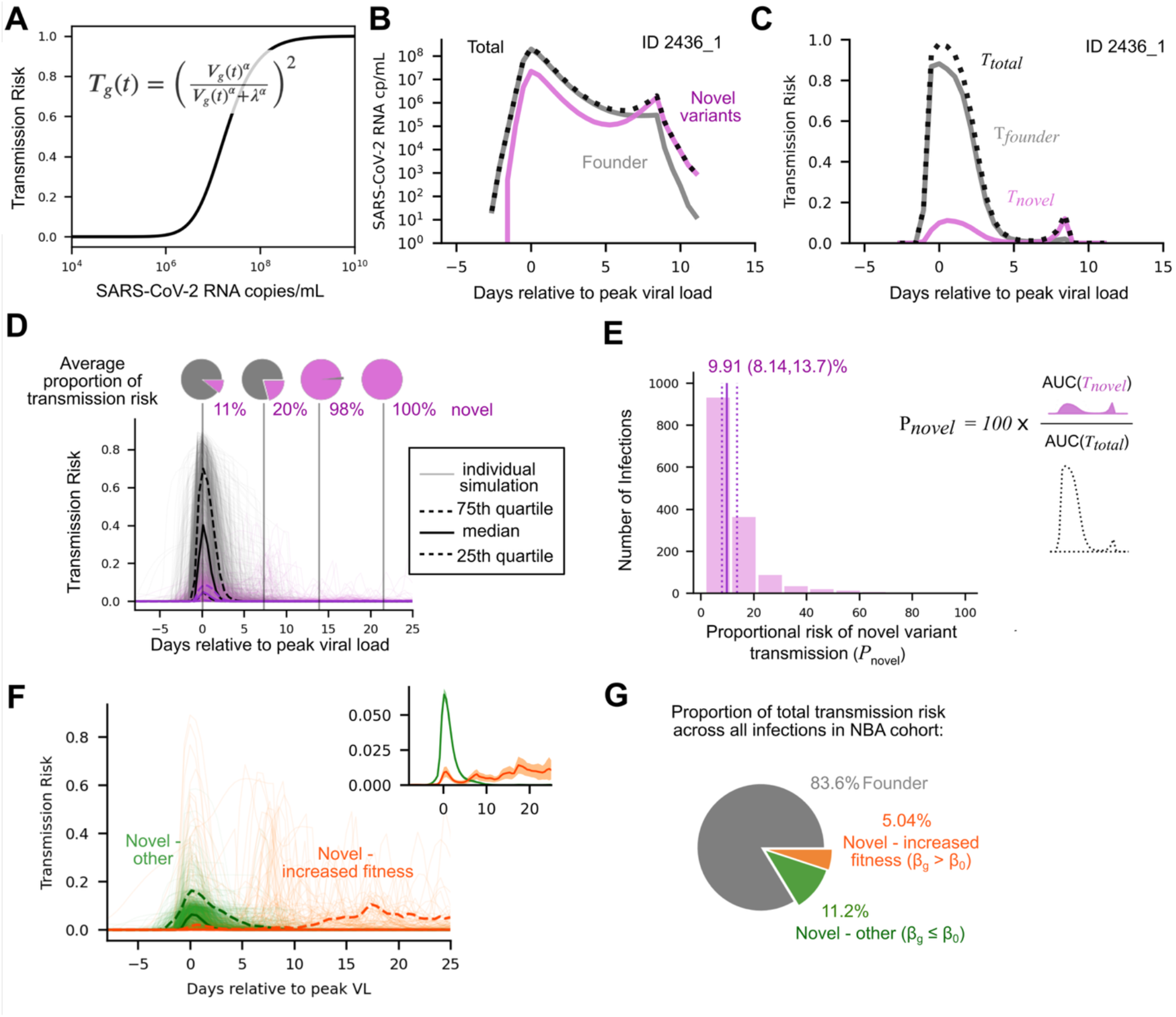
Dynamics of novel variant transmission risk. A) Data driven transmission risk model (**Eq 4**) relates viral load in copies/mL to transmission probability (0-1). 50% transmission risk of a given genotype variant occurs when V_g_ = λ. B) Example of viral load in one participant over the course of infection split by contribution to total viral load of founder vs. novel variants. C) Same participant as **B**, example of transmission risk attributable to founder vs. novel variants over the course of an infection. D) Transmission risk over time relative to peak viral load after simulating 1452 participants based on the NBA cohort. Thin lines show transmission risk of founder (gray) vs. novel variants (pink) with medians (bold) and IQRs (dashed lines) for both colors. Pie charts above represent mean risk of transmitting a novel variant given a transmission event occurs at peak viral load, and 1, 2 and 3 weeks post peak. E) Histogram of proportional risk of novel variant transmission during infection showing average 10% and IQR 8-14% in NBA cohort simulations. Inset shows calculation of proportional risk of novel variant transmission for the full infection: AUC of transmission risk for novel variants divided by AUC of total transmission risk x 100%. F) Transmission risk of novel variants over time relative to peak viral load after simulating 1452 participants based on the NBA cohort. Thin lines show transmission risk of novel variants with higher fitness than the founder variant (orange) vs. all other novel variants (green) with medians (bold) and 5^th^ and 95^th^ percentiles (dashed lines) for both colors. Inset shows the average transmission risk (bold) and SS% CI (shaded) with axis zoomed in to show transmission risk from 0-8%.

We applied the transmission risk model to WiPhy output from 1452 infection simulations representing the NBA cohort (**Fig 6D**). Variant takeover was relatively uncommon. In over 80% of infections the founder dominated throughout, and novel variants constituted an average of ~10% viruses at peak viral load and transmission risk. The total transmission risk decreased after peak, but proportion of risk attributable to novel variants grew from 11% at peak, to 20% a week after peak, and to 98% at 2 weeks **(Fig 6D)**.

The proportion of total transmission risk over the entire infection that was attributable to novel variants can be calculated as the area under the novel transmission risk curve divided by the area under the total transmission risk curve (**Fig 6E, inset**). Because total transmission risk was strongly driven by peak viral load, the proportion of the total transmission risk attributable to novel variants was on average approximately 10% (**Fig 6E**). Sensitivity analysis on the parameterization of the dose response model showed this result to be robust, varying from only 10-18% at parameter ranges across multiple orders of magnitude (**Fig S8**).

While the risk of transmitting any novel variant was highest at peak viral load, the dynamics of the transmission risk for novel variants with enhanced fitness were more nuanced. We further partitioned the novel variants between those with fitness-enhancing mutations (*β*_*g*_ > *β*_0_), and those without (*β*_*g*_ ≤ *β*_0_). Both the median and average risk of transmitting novel variants with fitness similar to the founder (S mutations) centers on the peak viral load **(Fig 6F, main panel and inset)**. However, the median risk of transmitting novel variants with increased fitness remained near zero throughout infection whereas the average risk rose around peak viral load and gradually increased as infection continued. Most infections had a small risk of transmitting a novel S variant at peak and negligible risk of transmitting a novel NS variant, but there were outlier cases that pose a substantial risk of transmitting a novel NS variant at peak.

We quantified the proportional risk of transmitting the founder versus novel variants with the same fitness as the founder versus novel variants with enhanced fitness across the NBA cohort by calculating the area under the respective transmission risk curves for each simulated infection (AUC of gray lines in panel D for founder, green lines in panel F for S variants, and orange lines in panel F for NS variants). Approximately 84% of transmission risk was attributable to the founder variant, 11% to novel variants with similar fitness to the founder, and only 5% to novel variants with enhanced fitness (**Fig 6G**).

### Prolonged infection as the most common source of transmission of a novel variant with enhanced fitness

While the degree of concordance between specific mutations which increased within host fitness and transmission fitness remains unknown, at least one new NS mutations is a pre-requisite to increase either type of fitness. Across the 1452 WiPhy simulations in the NBA cohort, we estimated that while 16% of the total transmission risk was attributable to novel variants, only 5% came from variants with a beneficial NS mutation (**Fig 6G**). During simulations, an average of one S mutation was detectable at peak viral load. In contrast, the average number of detectable NS mutations was zero until two weeks after peak (2.5-3 weeks into infection). Non-deleterious mutations that appeared early had a non-negligible chance of coexisting with the founder variant. Yet, due to the skewed fitness distributions (**Fig S2**), the probability of an emergent substantially beneficial NS mutation was more common after the viral population had time to expand and diversify. Beneficial mutations that did not appear in the first days of infection but ultimately fixed, required about a week to reach detectable levels, resulting in a further delay prior to detection or substantial transmission risk.

We categorized the infections simulated from the NBA cohort as founder-dominant, cleared takeover that occurred early (at least a week before infection cleared), cleared takeover that occurred late and posed negligible risk of NS transmission (<20% max), cleared takeover that occurred late and posed substantial risk of NS transmission, or takeover that resulted in persistent infection (duration > 40 days) (**Fig 7 inset**). Though founder-dominant cases constituted 80% of infections, they contributed only 7% of the overall risk of transmitting novel NS variants (**Fig 7 inset**). In contrast, cases of early variant takeover were rare, making up less than 4% of infections, but contributed 25% of the overall risk of transmitting novel NS variants (**Fig 7 inset**). The groups with an outsized contributions to the risk of transmitting novel NS variants were cleared-takeover that occurs late but when viral load was still high (1.5% of cases contributing 15% of risk) (**Fig 7 inset**) and persistent takeover cases (6.8% of cases contributing 50% of risk) (**Fig 7 inset**). In founder dominant, early takeover cases, and low-risk late takeover cases, the novel NS variant transmission risk was highest at peak viral load (**Fig 7A-C**). However, in the cases posing nearly 70% of the NS variant transmission risk, the risk was highest a week or more after peak viral load (**Fig 7D-E**). Overall, variants transmitted beyond a week post peak viral load were much more likely to be distinct from the founder variant, with an advantageous fitness phenotype.

**Fig. 7:**
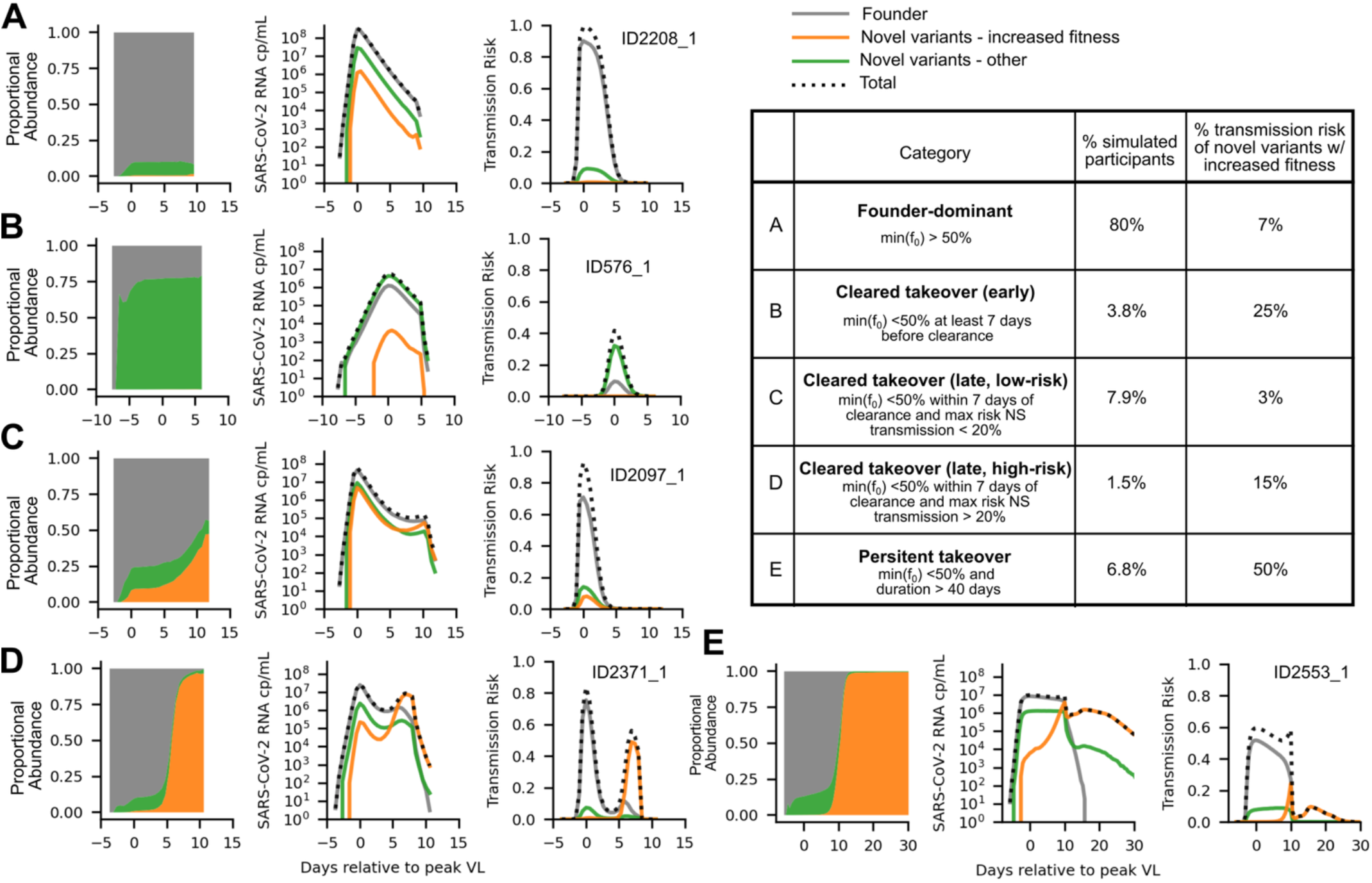
Examples of possible transmission risk profiles. The simulations of 1452 infections from the NBA cohort were classified as founder dominant, non-synonymous novel variant takeover cleared within 40 days, or non-synonymous novel variant takeover persistent for 40+ days for analysis. Cases of cleared takeover were further subdivided as early, late with a high risk of transmission, or late with a low risk of transmission. Here from left to right we illustrate the proportional abundance, the model predicted number of SARS-CoV-2 RNA copies/ml, and the predicted transmission risk of the founder variant (gray), novel variants with increased fitness such that β_g_ > β_0_ (orange), and other novel variants with β_0_ ≤ β_0_ (green). A) A typical example of a founder dominant dominant infection, which make up 80% of simulations (B-D) Three possible cases of cleared novel takeover. First, the novel variant emerges and predominates pre-peak (3.8% of infections), second the novel variant emerges late and predominates only briefly before infection is cleared (7.9% of infections), and third the novel variant emerges post peak but predominates while the viral load is high enough to pose a non-negligible transmission risk (15% of infections). E) A typical example of a persistent novel takeover infection. The time of takeover is similar to the onset of the acquired immune response, and the infection persists for the remainder of the simulation (6.8% of infections). Inset table summarizes how simulations were partitioned into these five takeover groups and what percentage of the total risk of transmitting a novel variant with enhanced infectivity within the NBA cohort comes from each group.

## Discussion

To examine how viral and immune dynamics within infected individuals shape SARS-CoV-2 evolution, we used a multiscale modeling approach. The mathematical model links viral characteristics, host-specific innate and acquired immune responses to project viral diversity, variant selection, and quasispecies fitness distributions. The model was validated against 16 phylodynamic metrics. We simulated a virtual cohort of ~1500 infections to identify specific viral fitness and immune conditions required for variant takeover. We projected the frequency of infections during which a variant was both fitter and achieved viral loads sufficient for transmission, a pre-condition for between-host spread. Cases of an early mutation conferring a fitness enhancement prior to peak viral load were extremely rare and occurred randomly. Two more predictable pathways to between host evolution were discovered: 1) delayed innate responses permitting rapid early viral diversification before adaptive responses control viremia, and 2) weak innate and adaptive immune responses permitting prolonged infections during which serially fitter variants accrue and persist at medium-high viral loads.

Our viral dynamic model has been validated against several natural history and clinical trial datasets^17–19^. Now, we extended it to phylodynamic data, matching the sampling depth and frequency of a household cohort study. Our simulations suggest that experimental divergence estimates correlate with and are only slightly lower than underlying model divergence. In contrast, we estimate only ~1% of circulating variants are detected with current experimental protocols but fortunately, most such undetected variants have deleterious or neutral mutations and have no lasting impact on viral fitness.

Our simulations project a highly skewed distribution of fitness among NS SARS-CoV-2 mutations which agrees with *in vitro* DMS data, further validating our fitting procedure. Throughout infection, the viral population includes a dense cloud of unfit, recently generated variants. The chance of a NS mutation increasing fitness by >1.5x was only ~0.3% and most NS variants with a fitness advantage relative to the current average fitness usually never predominated.

Beyond random mutation, dynamic innate and acquired immune responses drive variant selection. Because we do not assume explicit immune escape in variant fitness, if a variant with high relative fitness is expanding when immune forces are highly potent, then this variant will go extinct along with all other variants. Yet, moderate immune forces which do not eliminate the virus completely can enhance the natural selection of fitter variants by eliminating competing less fit viruses (purifying selection). Evidence of purifying selection has been found in numerous studies of within-host diversity^43–45^.

The assumption of broadly potent immune responses is based on substantial literature. For instance, innate immune forces follow type I interferon responses, which are not viral variant specific^46^. Acquired immune responses are more complex and naturally target specific viral epitopes, allowing the possibility of escape. But, the presence of redundant, overlapping clones of cross-reactive T cells^47,48^ suggests that immune evasion due to specific variant mutations may be relatively uncommon. Mutations that bypass humoral responses have been repeatedly documented and are relevant for loss of vaccine protection against infection^48–50^. It is less clear, however, whether endogenous antibody responses help contain SARS-CoV-2. Thus, we were unable to determine whether projected fitness increases due to new NS mutations are strictly due to enhanced cellular entry, or whether these mutations help bypass humoral and / or cellular immune responses. Our suspicion is that selection of specific mutations may occur for either or both reasons and is context dependent. The calibrated fitness landscape matches DMS data, suggesting viral variant intrinsic fitness may be enough to allow evolution. However, more detailed concurrent T cell and humoral response sampling will be valuable to further dissect this issue.

We next linked model-predicted host immune response features with viral evolution at the individual level. We found that certain immune parameter regimes favor founder-dominant infections with insufficient diversification and divergence for new variant predominance while other conditions favor novel variant takeover. Infections with a slower innate immune response defined by a low conversion rate of target to refractory cells, permitted a higher peak viral load and AUC that corresponded strongly with a higher maximum number of variants and HD. Yet, a delayed innate immune response was not sufficient for novel variant takeover.

Novel variant predominance was typically observed under parameter conditions when cells exited the refractory compartment at a moderate to high rate and there was a delayed or weak acquired immune response. Individuals with rapid cycling of target cells out of the refractory compartment developed a broader peak, allowing an extended period of viral diversification and divergence. This agrees with prior empirical observations that high intra-host diversity in early infection samples is predictive of prolonged infection^11,45^. This dynamic paired with an early but relatively weak acquired immune response favored persistent infection marked by extensive within-host evolution and expanding viral fitness. Fitter variant takeover in the absence of persistent infection was also a possible source of novel transmitted variants and occurred at moderate rates of refractory to target cell conversion.

We found different conditions favoring transmission of neutral versus fitter variants. The relative risk of transmitting a variant with an S mutation was ubiquitous across infections, accounting for ~10% of risk at the time of peak viral load. Viral AUC was the key determinant of transmission risk overall and thereby drove risk of transmitting a variant with a new S mutation. In contrast, beneficial NS mutations accounted for 1% of average transmission risk at peak viral load because beneficial mutations were sufficiently rare during the expansion phase to reach a meaningful quantity at peak viral load. In ~20% of infections, a variant with a beneficial NS mutations predominated after 10-14 days. Existing studies predict less frequent selective sweeps of new variants likely due to under sampling at these key late infection timepoints^51^.

Our simulations suggest that only ~10% of cases generate an NS variant at levels posing a transmission risk, suggesting a key bottleneck in the population viral mutation rate. Real-world contact patterns likely skew transmission events earlier in symptomatic infection^6,31,32,52–55^, which favors neutral mutations according to our simulations. Eighty percent of infected people do not infect another person. Superspreader events are required for a SARS-CoV-2 variant to outcompete other variants^35,36^. Given these multiple bottlenecks, and because population level sequencing likely misses most sub-dominant new variants, the observation of a single new branch on a phylogenetic tree is a remarkable outcome, contingent on the alignment of multiple immunologic and epidemiologic events.

The above findings are catered to transmission of a variant with only a single or small number of new base pair mutation. Most SARS-CoV-2 variants of concern have been highly mutated and emerged from ancestor viruses which had otherwise gone extinct in the population many months or years prior. The leading hypothesis for these saltation events is that highly immunocompromised hosts with persistent infections are a source. Our model suggests that transmission of a singly mutated virus likely could entail relative immunosuppression from the transmitter and that moderately prolonged infections are significant contributors to population level SARS-CoV-2 evolution. This prediction presents a public heath challenge because it suggests late (14+ days into infection) transmission events are rare but critical for viral evolution.

Our model has several limitations. First, we do not explicitly model recombination or epistatic interactions. Second, we do not map fitness changes at the level of single base pair mutations or gene. Instead, we calibrated the model to sequencing data from the spike coding region. Both of these additions are possible but would entail considerably greater computational complexity. Third, we only updated viral infectivity in response to mutations. The viral replication rate and other characteristics might also be subject to evolution^37,56^.Fourth, as discussed above, our model cannot discriminate variant specific immune evasion from enhanced cellular entry. Fifth, the immune mechanisms incorporated into the model also cannot be directly mapped to quantifiable cell populations or molecules but rather represent phenomenological immune pressure terms. Sixth, we assumed one well mixed viral population in the airways, but evidence of tissue compartmentalization and parallel evolution has been noted in case studies, mostly during persistent infection^9,57^. Seventh, our model generalizability might be somewhat limited by the different virus lineages and in vaccinated individuals^9,43,45^. However, our results suggest that differences in viral dynamics rather than mutation rates lead to different patterns of evolution across groups.

Finally, our model is not calibrated to data on infectious virus titers, so we assume relationships between measurements of the viral genome and transmission risk estimated in previous studies. Transmission risk is partitioned based on frequency of viral variants, making no distinction between within-host fitness and between-host fitness. For these reasons, the results of our transmission model are intended to be conceptual and quantitative model outputs are estimates.

In summary, we demonstrate deep links between host immune responses, viral evolution in a person, and spread of variants in the population. We conclude that rapid immune responses and short infection preclude selection of fitter variants during most infections, presenting a critical bottleneck of viral evolution rate in the population.

## Resource Availability

### Lead contact

Katherine Owens

### Data and code availability

Data and code used for the analysis presented in this work along with a more detailed derivation of the within-host effective reproductive number can be accessed at https://github.com/lacyk3/SARS-CoV-2WithinHostEvolution.

## Acknowledgements

This research was funded by awards from the National Institutes of Health (NIH) National Institute of Allergy and Infectious Disease including K25AI196259 to KO and R01AI186721 to DBR. This research was supported in part by the Intramural Research Program of the NIH. The contributions of the NIH authors, PR and EB, are considered Works of the United States Government. The findings and conclusions presented in this paper are those of the authors and do not necessarily reflect the views of the NIH or the U.S. Department of Health and Human Services.

## Author Contributions

Conceptualization: KO, DBR and JTS with consultation from PR, ST, AG, CC and EB

Methodology: KO and DBR

Software: KO and DBR

Validation: KO

Formal Analysis: KO

Visualization: KO

Data Curation: PR and EB

Original Draft: KO

Review & Editing: KO, DBR, JTS, PR, ST, AG, and CC

## Declaration of Generative AI and AI-Assisted Technologies

This work is solely created by the author(s). No generative AI or AI-assisted technologies were used.

## Methods

### Experimental Model and Study Participant Details

For model development and calibration, we relied on paired viral load and sequencing data from longitudinal upper respiratory specimens collected by Cohen et al. from two cohorts in South Africa between October 2, 2020 and September 30, 2021^38^ or between May 1, 2020 and December 31, 2020^39^. The former constitutes a case-ascertained household infection cohort and the latter a hospitalized cohort. Ko et al. then performed high-throughput, single-genome amplification and sequencing (HT-SGS) on the SARS-CoV-2 spike gene from each sample. By considering the unique combinations of called mutations (i.e., haplotypes) that are supported by multiple single-genome sequences (SGS), HT-SGS demonstrates mutational linkage patterns across the 3.8-kilobase spike region that are not detectable by short-read whole-genome sequencing. Using the spike sequences in each sample, Ko et al. calculated statistics that summarize within host diversity including the number of detectable haplotypes present, the average pairwise distance between two sequences, and the frequency of the first dominant detected haplotype. Ko et al. calculated the average pairwise distance for each group of sequences from a single sample as the total number of mutations (point mutations and/or indels) between each pair of sequences divided by the number of possible pairs in that group. In our results and discussion, we refer to variants instead of haplotypes. Further detail on the sequencing technique used can be found in Ko et al ^11^.

We filtered the data analyzed by Ko et al. to infections that cleared within 40 days of symptom onset, were documented with at least 3 quantitative viral load measurements, and did not occur in people with uncontrolled HIV. This included N=14 infections in people without HIV and N=4 infections in people living with HIV with CD4 T cell count ≥ 200 cells/µL. We chose to include individuals living with controlled HIV when fitting the WiPhy model because Ko et al. reported that these two groups exhibited similar intrahost diversity and viral dynamics^11^. In this group of 18 infections, viral load was quantified at a median of 4 (min 3, max 13) timepoints and a median of 3 (min 1, max 8) timepoints had sufficiently high viral load for sequencing. Sampling started a median of 4 (min −3, max 17) days post symptom onset and occurred every second day or three times weekly until viral clearance.

Information on biological sex was collected in these studies through self-reporting. While sex was considered as a potential exposure of interest in the design of the original studies, analysis was not stratified by sex in the current study.

### Within-host Phylodynamic Model

In the absence of mutation, the mathematical model is the stochastic analog of our previously developed and data-validated model for acute SARS-CoV-2 infection^17^ **(Fig 1A)**. The system describes cells that are susceptible to SARS-CoV-2 infection, denoted *S*, which become infected upon interaction with virus, *V*, with rate *β*. Infected cells initially enter an eclipse state, *I*_*1*_, which they remain in for an average time 1/*k* before transitioning to a productively infected state, *I*_*2*_. Productively infected cells output virus at an average rate *π* for an average lifespan of 1/*δ* days prior to the onset of the acquired immune response. The presence of productively infected cells triggers innate immune signaling that pushes the susceptible cell population into a state that is refractory to infection, *R*, at an average rate *ϕI*_2_. Refractory cells revert to a susceptible state at an average rate of *ρ*. At *τ* days post infection, an adaptive immune response emerges which increases productively infected cell clearance rate to *δ* + *m*.

The rules governing the mechanistic model inclusive of mutation can be expressed as a set of stochastic differential equations describing a branching process. As novel genotypes (indexed with parenthetical superscript integers [1,*N*_*t*_]) are generated, the number of infected cell and viral compartments, *N*_*t*_, grows. At time *t* the dynamics of infection are given by Eq. (1)

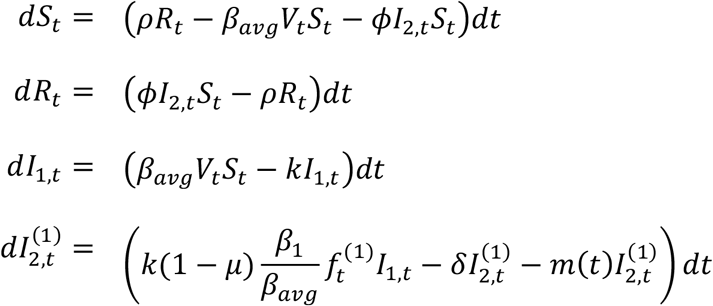

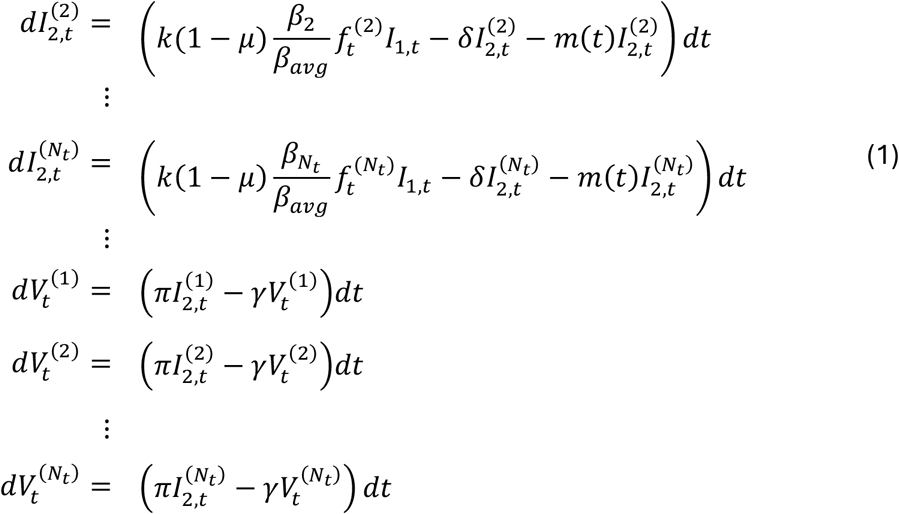

Where 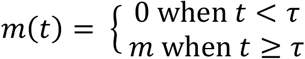 is a step function that implements a delayed acquired immune response, 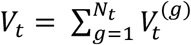 is the total viral population, 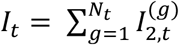 is the total productively infected cell population, and the current average infectivity of the viral population is

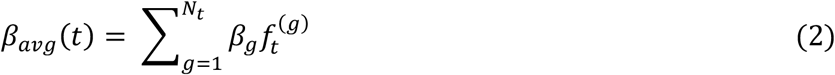

where 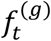 denotes the frequency of variant g at time t, 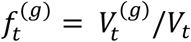.

The force of infection of the total viral population,

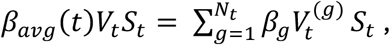

determines the number of newly infected cells entering the eclipse state. The proportion of infected cells transitioning from the eclipse to the productive state that are infected with variant *g*′ is determined by the frequency and relative infectivity of *g*′ at time t, 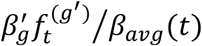. On average (1 − *μ*) of the cells newly infected with variant *g*′ will produce variants of the same genotype, but with probability *μ* the infected cell is the first of a new variant, *g*′′, that has *g*′ as its parent. This expands the number of compartments, so *N*_*t*+1_ ≥ *N*_*t*_.

### Details of WiPhy Model Implementation

The model is implemented in Python using a discrete stochastic τ-leaping simulation scheme calibrated to data in 1 mL of fluid and a simulation time interval of Δt = 0.01 days. The state variables,

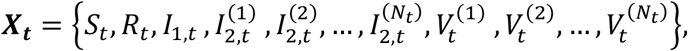

represent the numbers of susceptible, refractory, eclipse and productive infected cells and virions (for each genotype) respectively. In each time interval, a Poisson-distributed (℘) number of events, ***e***, of each mechanistic transition is chosen using a rate parameter equal to the product of the reaction propensity, *p*(***X***_***t***_), and the size of the timestep such that ***e***~ ℘(*p*(***X***_***t***_)Δ*t*). The possible transition events are the terms on the right-hand side of system 1, like cell death, infection of a new cell, viral production or clearance, etc.

After the number of transition events has been determined, the possibility of mutation is incorporated as cells transition to the productively infected state. Each newly productive infected cell is assigned a genotype based on one or more binomial draws. Infected cells of a given genotype produce virions of the same genotype for their whole lifespan. The first draw determines whether the infected cell will produce virions that are genetically distinct from its infecting (parent) virion with probability *μ* or produce virions identical to its parent virion with probability 1 − *μ*. If the cell is producing identical virions, then the total number of cells of the parental variant is increased by one. If the cell is producing genetically distinct virions, it is assigned a new integer genotype *g*, and a second binomial draw determines whether the new genotype has the same phenotype or a distinct phenotype compared to the parent virion with probabilities *η* and 1 − *η* respectively. Novel genotypes with the same phenotype as their parent are assigned the same infectivity, *β*_*g*_ = *β*_*parent*_, and their genotypic change is assumed to be a single synonymous mutation. Novel genotypes with a distinct phenotype are assigned a new infectivity by drawing a scale factor from an exponential distribution, Δ*β*_*g*_~ *Exp*(⟨Δ*β*⟩), and multiplying this fitness-effect with the infectivity of the parental variant such that *β*_*g*_ = Δ*β*_*g*_ × *β*_*parent*_. The number of nucleotide changes associated with this mutation event is drawn from a Poisson distribution Δ*hd*_*g*_~ ℘(⟨Δ*hd*⟩). Allowing Δ*hd*_*g*_ > 1 allows for the possibility of non-point mutations, like small indels or multinucleotide mutations.

Once the number of non-mutated vs. mutated infection events has been determined, and ***e*** adjusted accordingly, the state variables are updated using the event transition matrix, *T*, so at the next time step for existing variants ***X***_***t***+**Δ*t***_ = ***X***_***t***_ + ***e****T*. Then the list of state variables is extended to include the newly created variants.

The initial condition used for each simulation matched the initial conditions used to estimate viral dynamic parameters for the ODE model for the NBA cohort. However, we initialized the number of infected cells in the eclipse stage at two in order to reduce the number of simulations that stochastically burn out early, prior to reaching detectable viral levels. This led to 1452 usable simulations out of 1510 runs in the NBA cohort (98%).

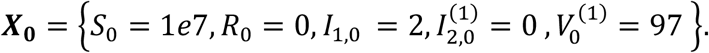

### Targets for Parameter Fitting

We defined 16 quantitative metrics of interest and calculated their distribution in the SAI cohort, illustrated in **Fig S1A**. To summarize viral dynamics in the cohort, we calculated the peak viral load, the time to peak viral load, the time near (within 1 log) peak viral load, and the duration of infection in days. To summarize viral diversity, we calculated the number of unique variants detected and the average pairwise genetic distance at days 2, 5, 10 and 20 post symptom onset. We also aimed to match the waning of the founder virus by calculating the detected frequency of the first dominant variant at days 2, 5, 10 and 20 post symptom onset. See **Table S1** for definitions, calculation, and mean and variance of data targets.

### Details of Parameter Fitting Scheme

By using previous estimates from the literature, we reduced parameter estimation to 3/10 remaining parameters. **Table S2** provides information on fixed parameters and estimated parameter ranges. This scheme assumes some intrinsic viral characteristics are shared across all infections.

We fixed the fraction of S mutations at *f*_*s*_ = 0.35 based on previous estimates in the literature^40,41^. Then, we applied a grid search over the parameters governing NS mutation, covering the domain

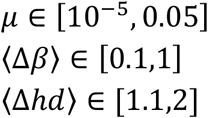

For each tuple consisting of a candidate mutation rate, *μ*, expected change in fitness given NS mutation, ⟨Δ*β*⟩, and expected change in hamming distance given NS mutation, ⟨Δ*hd*⟩, we drew 30 viral dynamic parameter sets from those previously estimated to describe infections in the National Basketball Association cohort. We ran these 30 simulations with the candidate evolutionary parameters and saved the output every 12 hours. We randomly down sampled each WiPhy simulation run as described in the following section, *Down-sampling Simulation Output for Comparison with Data*, to obtain coarse-grained simulation output. If a simulation did not result in detectable infection due to stochastic burn out, it was left out of subsequent analysis because such individuals would not have been included in the SAI cohort.

We calculated the distribution of the target metrics illustrated in **Fig S1A** and reported in **Table S1** using the coarse-grained model output. Then, we applied the two-sided Kolmogorov–Smirnov (K-S) test for the equality of the distribution calculated from SAI data and from WiPhy output for each metric. If the test returned a p-value of p>0.05, indicating that the distributions were not considered different in a statistically significant manner, the parameter set scored a point for that metric for a maximum of 16/16 points (**Fig S1B**). We repeated the down sampling and scoring process 10 times and used the average score across these 10 instances of sampling to determine the best fit parameter values.

### Down-sampling Simulation Output for Comparison with Experimental Data

To approximate the experimental procedure that generated sequence data metrics (Ko et al.^11^ in the SAI cohort) we applied a coarse-grained sequence sampling. Simulated viral load was used to choose a three-category sequence sample size. If simulated SARS-CoV-2 RNA copies/ml V<10^4.25^, no sequences were recovered (n=0), if 10^4.25^<V<10^6^, n=10 sequences were recovered, and if 10^6^<V, n=500 sequences were recovered. **Fig S2B** illustrates how this scheme compares to the empirical distribution of SGS recovered in the SAI cohort data. Sequence selection was achieved by randomly choosing n sequences (infected cells) from the simulated cell population and counting sequence variants in this sample. We then enforced that at least 2 copies of a sequence must be present for that variant to be detected, and a minimum detectable frequency per sample was 0.45%. These two criteria respectively excluded sequences appearing in less than 2/10 sequences or 3/500 sequence sample. At higher viral loads, this is only slightly more sensitive than Ko et al.’s protocol, which required 5/500 sequences^11^.

For each simulated infection, we also down sampled in time when evaluating agreement with the SAI cohort data. An incubation period was drawn from the distribution of known incubation periods in the NBA cohort, **Fig S2D**, and a random delay between symptom onset and the time to first sample was drawn from a uniform distribution with a median of 4 days (IQR 3-8) after symptom onset. Once the day of first sample was determined, we sampled the simulation every 2 days for the first 14 days, then reduced the sampling frequency to every 5 days for the remainder of the infection if applicable.

### Model Validation via Individual Simulations

We first validated the evolutionary parameter estimates by assessing whether the population level estimates were able to reproduce individual level data. We estimated individual viral dynamic parameters for each infection documented in the SAI cohort using a population non-linear mixed effects approach implemented in Monolix. This approach models each viral load measurement from an individual *i* at a time point *k* as log_10_(*y*_*ik*_) = *f*_*V*_(*t*_*ik*_, *θ*_*i*_) + *ϵ*, where *f*_*V*_ represents the solution of the ODE model for the state variable describing the virus, *θ*_*i*_ is the parameter vector for individual *i*, and *ϵ* ~ *N*(0, *σ*^2^) is the measurement error for the log10-transformed viral load data. Furthermore, in the population model, each individual’s parameters can be written as the sum of the average population value *θ*_*pop*_ and a random effect encompassing their deviation from the average, *η*_*i*_; the parameters for individual *i* are given by *θ*_*i*_ = *θ*_*pop*_ + *η*_*i*_. The parameter estimates are available in the supplementary material.

Then, we ran 10 WiPhy simulations using each of these viral dynamic parameter sets along with the population estimates for the evolutionary parameters obtained from the grid search. We randomly down sampled the variants produced by each WiPhy simulation as described in the previous section, *Down-sampling Simulation Output for Comparison with Data*, 10 times independently. We compared each instance of sampling a trajectory against the individual level data and identified the best fit with regards to the sum of squared error for viral load, number of variants detected, average pairwise distance, and frequency of the first dominant variant. Then we picked the “best” run/sample combination as that which had the lowest ordinal ranking across the four metrics. In the event of a tie, we picked via visual inspection. Three examples of the individual-level agreement are shown in the main text in **Fig 3** and all individuals are shown in **Fig S3**.

### Constructing a Family Tree of Detected Sequences

For all infections in the SAI cohort and a subset of the infections in the NBA cohort, we constructed trees depicting the family lineage of variants that reached a threshold maximum frequency over the course of infection. WiPhy simulation output lists the parent variant from which each novel variant emerged via mutation. When we reduce the number of variants to a “detectable” subset of the list, we check which detectable variant is the most recent ancestor of each variant and save this information. This can be translated into a directed adjacency matrix, *A*, with detected variants arranged along the rows and columns ordered by birthdate and a non-zero entry at *a*_*i,j*_ if variant *i* is the most recent ancestor of variant *j* among the set of detected variants. To visualize the resulting trees we used the NetworkX package in Python^58^. Each detectable variant in the subset of interest is represented with a node and an edge connects the variant back to its most recent ancestor. The trees are always rooted at the founder variant. In the visualizations included in the manuscript, we consistently determine the horizontal position of each node from the variant birthdate, the color from the HD from the founder variant, and the radius from the log of the number of progeny variants.

### Transmission Models

We predicted SARS-CoV-2 transmission risk using a dose-response curve that maps from viral RNA copies/mL to risk of transmission to exposure contacts, i.e. those who are physically exposed to the virus. Goyal et al. calibrated this model using individual reproduction number^59^ and serial interval data^60,61^ when studying SARS-CoV-2 superspreading. Goyal et al. posit that only some exposure contacts have virus passed to their airways (contagiousness) and only some exposed contacts with virus in their airways become secondarily infected (infectiousness). Both infectiousness and contagiousness are calculated from viral load using the hill function,

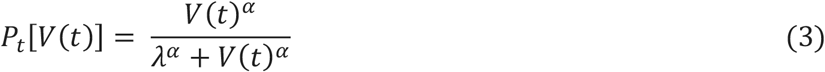

Here, *λ* is the infectivity parameter that represents the viral load that corresponds to 50% infectiousness and 50% contagiousness, and *α* is the Hill coefficient that controls the slope of the dose-response curve. Their calibrated model was further validated by equation 3 with the optimal parameter values with in vitro probability of positive virus culture from van Kampen et al.^62^ and finding good agreement.

Contagiousness and infectiousness are then treated as viral load dependent multiplicative probabilities with transmission risk for a single exposure contact being the product,

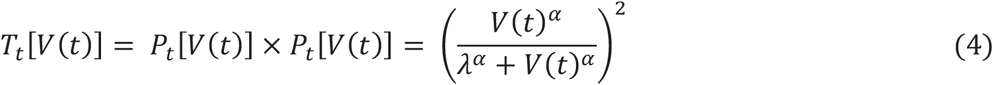

We partitioned the total transmission risk into variant-specific transmission risks using the frequency of each variant over time.

### Sensitivity Analysis on Transmission Risk Parameters

The use of dose-response curves to map viral load to infectious virus is widespread^23,24,63,64^ though the specific functional form used varies and optimal parameter estimates differ depending on the calibration data set. We explored the robustness of our conclusions regarding transmission risk to variability in the transmission risk parameters. The dose-response curve is shown for varying viral load at 50% contagiousness *λ* ∈ [1*e*5, 1*e*9] and varying hill exponent *α* ∈ [0.2,2] in **Fig S8Ai-iii**. We quantified the total risk of transmission that the infection poses by calculating the area under the transmission risk curve (although this assumes there are not drastic changes in behavior). The percentage of total transmission risk attributable to novel variants is then taken to be the area under the novel transmission risk curve over the area under the total transmission risk curve times 100 (**Fig S8Aiv-v)**.

We applied the transmission risk model with combinations of *λ* ∈ [1*e*5, 1*e*9] and *α* ∈ [0.2,2] to the WiPhy output from the 1452 infections in the NBA cohort. Then, we calculated the average transmission risk over the course of infection under each dose response curve and visualized the outcome as a heatmap (**Fig S8B)**. As expected from the red curves in **Fig S8A**, the risk of transmission over the course of infection is much higher for lower values of *λ* and higher values of *α*. However, the average proportion of transmission risk attributable to novel variants, shown in **Fig S8C**, only ranges from 10-18%. We then aligned the curves temporally based on the timing of peak viral load and considered how the variation in transmission risk over time depends on *λ* when the hill exponent is fixed at *α* = 1 (**Fig S8D**). Regardless of *λ*, the transmission risk is always highest at peak viral load. For lower values of *λ*, the transmission risk remains elevated for a few days after peak viral load and for *λ* = 1*e*5 the average transmission risk also picks up again due to viral rebound 20 days after the peak. The proportion of transmission risk attributable to novel variants at a given moment in time is independent of *λ*, with the proportion of risk always rising as the infection continues (**Fig S8E**).

### Converting Ct Measurements into Viral Load

Meiring et al.^39^ and Kleynhans et al. performed RT-PCR using the Allplex™ nCoV 2019 kit (Seegene, Seoul, South Korea). Specimens were considered positive for SARS-CoV-2 nucleic acids if the Ct was < 40 for ≥ 1 of 3 gene targets. Though all specimens were tested in the same laboratory, using standardized technique and assays, they stopped at this qualitative assessment of viral load rather than generating calibration curves to determine quantitative viral load. We converted cycle threshold (Ct) values for the N gene collected from the SAI cohort to viral genome equivalents, using the same equation employed in our previous modeling work in Owens et al.^17^

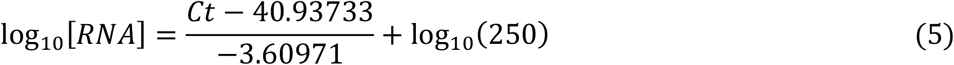

This equation was reported by Kissler et al.^3^, who estimated the parameters using specimens across a 13-point standard curve (dynamic range from 250 copies/mL to 4.50 × 10^8^ copies/mL). The specimens were generated at Caltech and underwent extraction using the FDA-authorized Quick SARS-CoV-2 RT-qPCR. There is close agreement between this equation and the standard curve for the Seegene Allplex assay reported by Jiminez et al:

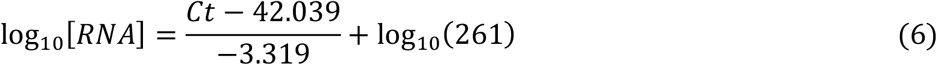

We noted that the difference between these two curves across the dynamic range 250 copies/mL to 4.50 × 10^8^ copies/mL is always less than 1 on a log scale, which is close to the model predicted measurement error for viral load. Opting to use the conversion equation calibrated by Kissler et al. simplified the computational pipeline.

### Calculating Average Pairwise Distance from WiPhy Simulation Output

The WiPhy model does not track nucleotide sequences, so to calculate the number of base-pair differences between a pair of sequences we compute the sum of their HDs and subtract off the HD of the sequence of their most recent common ancestor Δ(*i, j*) = *H*_*i*_ + *H*_*j*_ − 2H_P(i,j)_. The most recent common ancestor is determined by comparing the parents of both variants. If the parent is identical, the procedure halts and the parent is considered H_P(i,j)_. If the parent is different, we track back to parents of the parental sequences and repeat. We record the maximum HD from the founder sequence 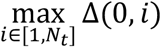 as a measure of divergence. Here variant 0 is the founder sequence. We calculate the average pairwise distance among, *S*_*t*_, the unique set of sequences present in a sample from time *t*, as:

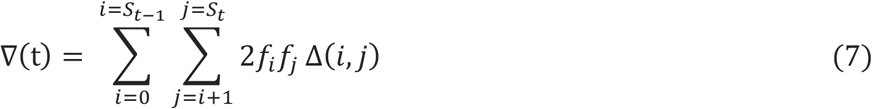

where *f*_*i*_ is the detected frequency of the *ith* unique variant in the sample. This allows comparison with the value from Ko et al., who calculated average pairwise distance for each group of sequences as the total number of mutations (point mutations and/or indels) between each pair of sequences divided by the number of pairs in that group.

### Calculating the Effective Within-host Reproductive Ratio

We used the next-generation method developed by van den Driessche and Watmough^65^ to determine the within-host basic reproduction number of SARS-CoV-2, ℛ_0_. The within-host basic reproduction number is the expected number of secondarily infected cells produced in a completely susceptible population by the introduction of a single “typical” infected cell^66^. In a single-variant model without evolution the reproductive number is ℛ_0_ = *S*_0_*β*_0_*π*/*γδ*_0_, which is the same expression as a basic target-cell limited model without an eclipse or refractory compartment.

The reproductive number of a novel variant that emerges during infection depends on the effective reproductive number, ℛ_*t*_, at the time that it is generated and its fitness relative to the current population average. Consider one genotype of interest, *g*, and group all other existing genotypes into one viral compartment. The condition under which virus with genotype *g*is expected to increase in frequency if it emerges at time *t* is (1 − *μ*)*β*_*g*_ > *β*_*avg*_(*t*) where *β*_*avg*_(*t*) is the average infectivity of the existing variants weighted by their current frequencies, *β*_*avg*_(*t*) = ∑_*i*≠*g*_ *β*_*i*_*f*_*i*_(*t*). Furthermore, the effective reproductive number of genotype, *g*, must be greater than one for the expected population of cells infected with variant *g* to be growing. That is, 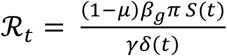 must be greater than one. Note a variant could grow in frequency even as its population is shrinking in size if it is shrinking more slowly than the rest of the population. Conversely, a variant’s population could grow even as its frequency decreases if it is growing more slowly than the population average. Furthermore, these conditions govern the expected behavior of an emergent genotype, but in each iteration of the stochastic process when the size of the cell populations are small random fluctuations may lead to extinction of variants even with ℛ_*t*_ > 1. For a derivation of these results please see the supplementary file on github at https://github.com/lacyk3/SARS-CoV-2WithinHostEvolution.

