## Supplemental Figures and Tables for "Transmission of mutated SARS-CoV-2 variants is favored by relatively prolonged infections due to delayed immunity"

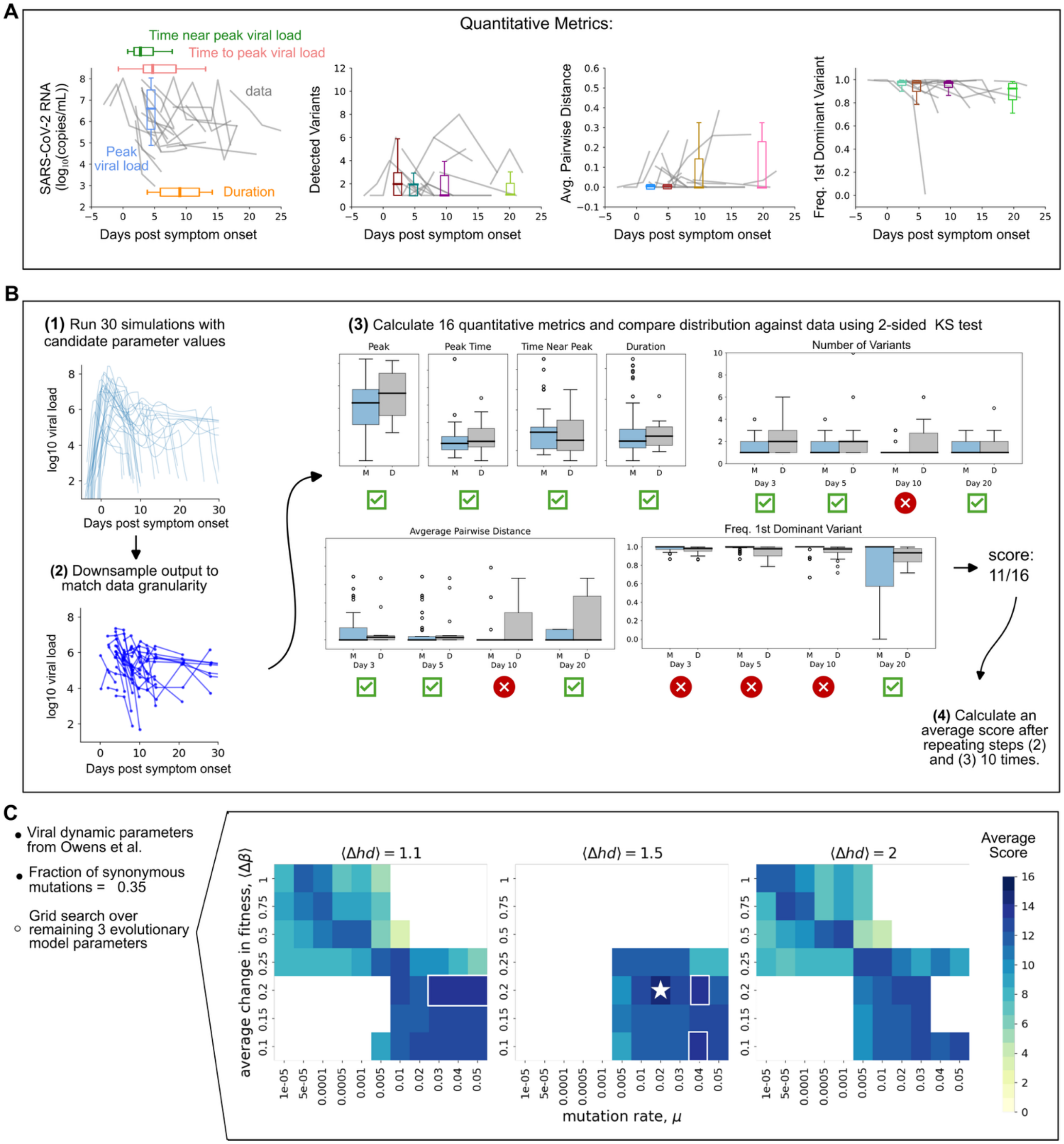


***Fig S1. Model Calibration Approach.***

*A) Data trajectories (lines) and model fitting metrics including number of detected variants, average pairwise distance and frequency of the first predominant strain (boxes). Metrics associated with values and timing are shown as vertical and horizontal boxes, respectively. All boxplots are median, interquartile range (IQR, box) and 1.5x the IQR (whiskers).*

*B) The model fitting workflow follows 4 steps: (1) A full run involves choosing a single parameter set including mutation rate $\mu$, average change in fitness $\langle \Delta\beta \rangle$, and average change in Hamming distance $\langle \Delta HD \rangle$ and outputting a finely simulated viral load trajectory and associated sequence evolution metrics. Here 30 runs are shown with different viral dynamic parameter values drawn from the population values estimated for the NBA cohort in Owens et al. (2) The simulation output is sampled to match the experimental design of data collection for the SAI cohort in terms of frequency of sampling (1-7 days between samples) and the depth of sequence sampling (See Methods and Supplementary Figure 2). (3) From these output data, kinetic and* *evolutionary metrics are calculated for direct comparison between model and data. The "score" of the given model parameter set is computed as the number of metrics for which data and model are similar (as evaluated by the two sided KS test for similarity of distributions between the data (gray) and model values (blue)). All boxplots are median, interquartile range (IQR, box) and 1.5x the IQR (whiskers). In the example shown here, the model run with these parameters produces a poor fit to the data resulting in a low score of only 11/16.*

*C) Across the three calibrated evolutionary model parameter values, a combination of $\mu$=0.02, $\langle \Delta\beta \rangle$=0.2, and $\langle \Delta hd \rangle$=1.5 resulted in an average score of 16/16 for 10 independent instances of down-sampling the simulation output and scoring (starred). 5 other parameter sets averaged a score of 15/16 for 10 instances of down-sampling (boxed).*

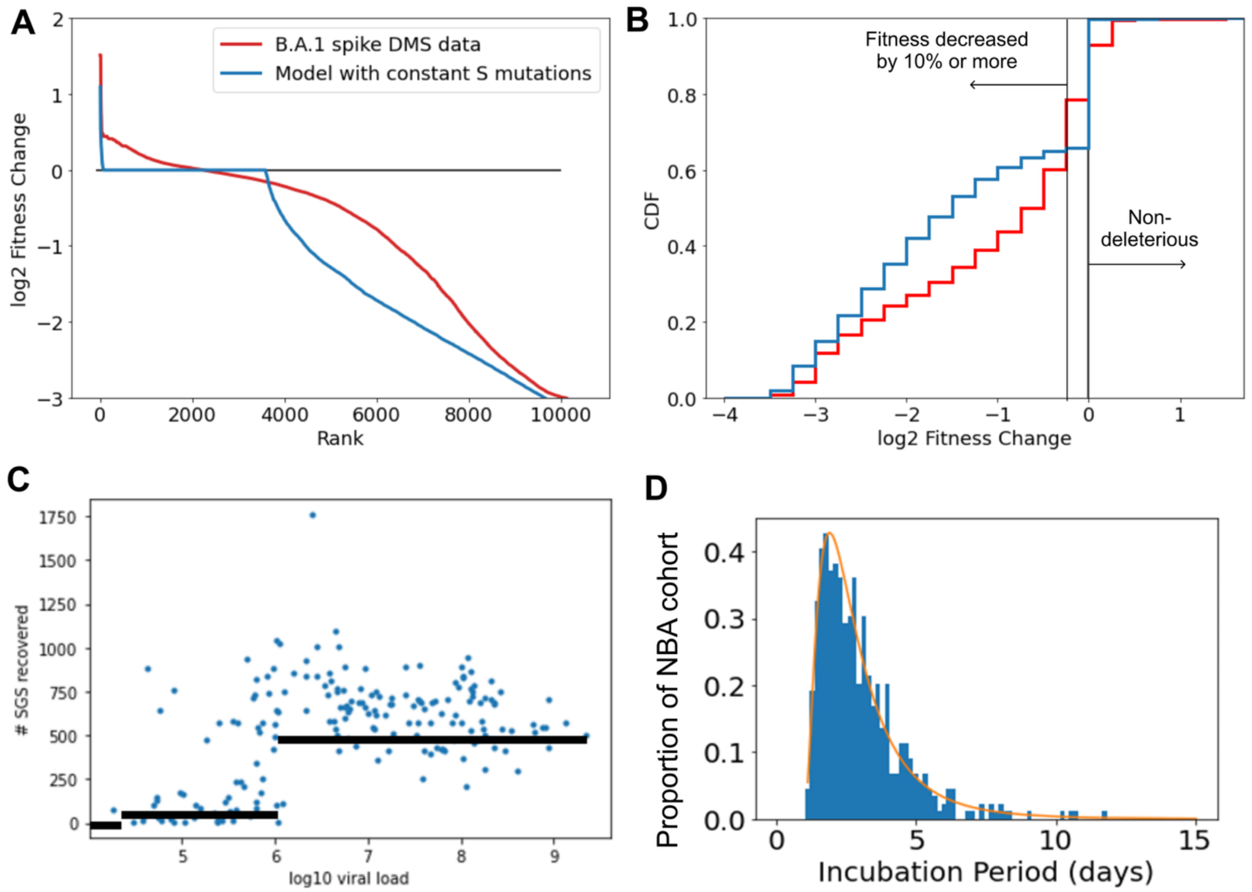


***Fig S2: Comparing WiPhy Model with Data.***

*A) Rank-fitness effect curve from the calibrated evolutionary model in blue compared with the experimental rank-fitness effect curve for mutations within the spike region generated by Bloom et al. in red. The model assumes that all synonymous mutations have an identical fitness resulting in a broad horizontal line at fitness change of 0 on a log scale. The experimental data intersects 0 within this region.*

*B) Cumulative distribution function for fitness effect distribution with the model in blue and experimental data in red. The fraction of mutations predicted to be neutral or beneficial by the calibrated model is 35.5% compared with 21.3% neutral or beneficial mutations observed in the data.*

*C) Each blue dot is a sample sequenced by Ko et al. plotting the number of single-genome sequences (SGS) recovered versus the viral load (SARS-CoV-2 RNA cp/mL) of the sample. Black lines indicate the viral load-dependent number of sequences drawn from the WiPhy simulation output when down sampling with respect to viral diversity to calculate detectable quantities.*

D) The distribution of incubation periods estimated based on ODE model fit to infections with a known symptom onset in the NBA cohort. This distribution was used to draw an incubation period to align trajectories based on symptom onset and compare with SAI data.

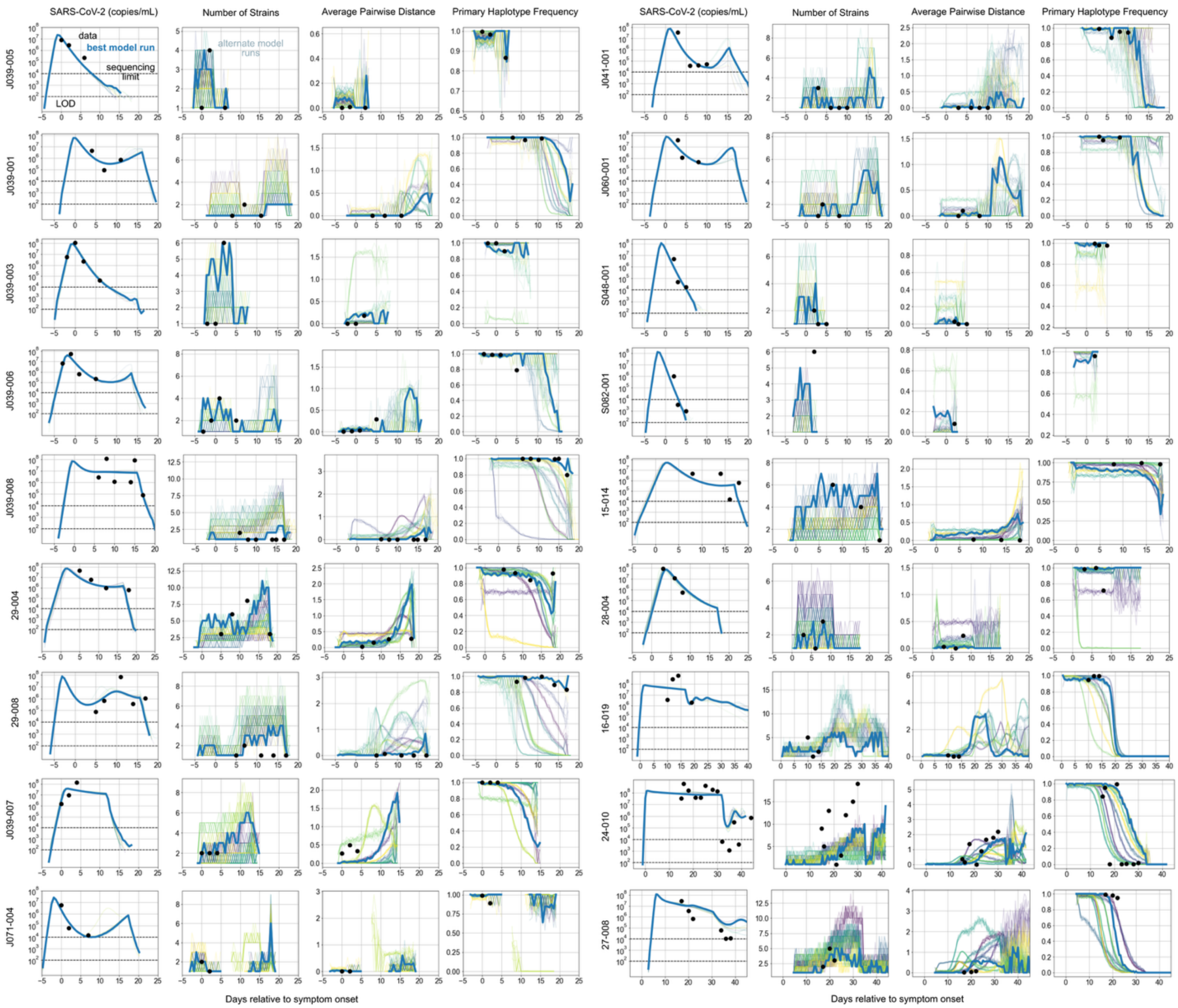


**Fig S3.** ***Individual phylodynamic output for SAI participants generated using population-calibrated model.*** *Comparison of model (lines) and data (dots) from 18 participants in the SAI cohort with acute, self-limited viral infections and viral load quantified at a minimum of 3 data points. 10 replicate simulations (colors) with 10 each stochastic subsamples (thin lines of same color) with bold blue line highlighting the sampled trajectory that best matched the four metrics. From right to left: viral load, number of detectable variants, average pairwise genetic diversity of detected variants, and the frequency of the first dominant variant. Viral dynamic model parameters in each example were chosen by best fit to viral load data and the population calibrated evolutionary model parameters,* $\mu$*=0.02,* $\langle\Delta\beta\rangle$*=0.2, and* $\langle\Delta hd\rangle$*=1.5, fs = 0.35, were used.*

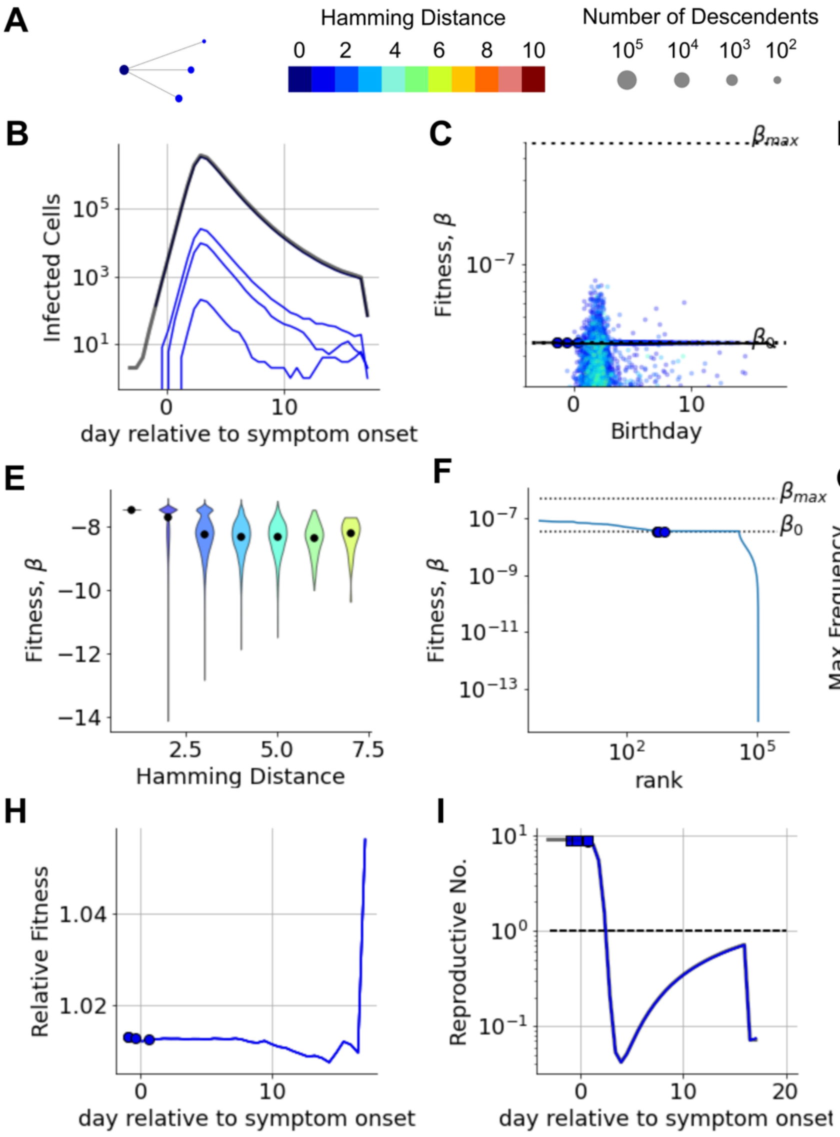


***Fig. S4: A detailed look at fitness evolution during a typical acute infection.*** *Panels A-I were generated using model output from one WiPhy simulation using population-calibrated evolutionary parameters and viral dynamic parameters estimated based on data from infection in one individual.*

*A) Simulated lineage of major variants (those reaching ≥2% of viral load during infection). Node radius is proportional to number of descendants (log10) and horizontal position corresponds to variant birthday (first time it appeared in simulation relative to symptom onset) concurrent with panel B. This color bar indicates HD for all panels* ***A-J****.*

*B) Simulated number of productive infected cells ($I_2$) corresponding to variants in* ***A*** *and their total (bold black).*

*C) Relationship between variant birthday and absolute variant fitness. Single variants (small dots colored by HD) show heterogeneity and mean (black line) trails the fitness of emerging major variants (larger dots). Dashed horizontal lines denote initial variant fitness $\beta_0$ and maximum $\beta_{max}$ fitness allowed in the simulation.*

*D) Violin plots of fitness of emerging variants grouped by 5 day intervals indicate bimodal fitness distributions with growing difference between mean (black line) and median (black dot). Note linear scale in comparison to* ***C****.*

*E) The bimodal fitness distributions of variants grouped by HD show increasing variance as HD increases. Median value indicated by black dot.*

*F) Each variant's fitness plotted by rank (ordered from highest to lowest infectivity). Line shows complete distribution with large dots indicating major variants. Horizontal lines following major variants reflect synonymous variants. Many variants with fitness higher than the founder never reach detectable levels.*

*G) Maximum frequency reached by variants versus their fitness relative to the population mean at time of variant appearance is roughly log log linear. Large dots indicate major variants above 2% frequency. These and other detectable minor variants (frequency above the limit of detection, LOD) have relative fitness near or above 1.*

*H) Time dynamics of the relative fitness of major variants showing that relative fitness is highest when a strain emerges and then decreases – note linear scale in comparison to* ***E*** *in which many variants have small but heterogeneous fitness.*

*I) Effective reproductive number (average number of newly infected cells from one infected cell,* ***Methods****) for each major variant (dot at birthday with line showing dynamics) compared to the population average (black line) over time after symptoms. New variants only can predominate when R>1.*

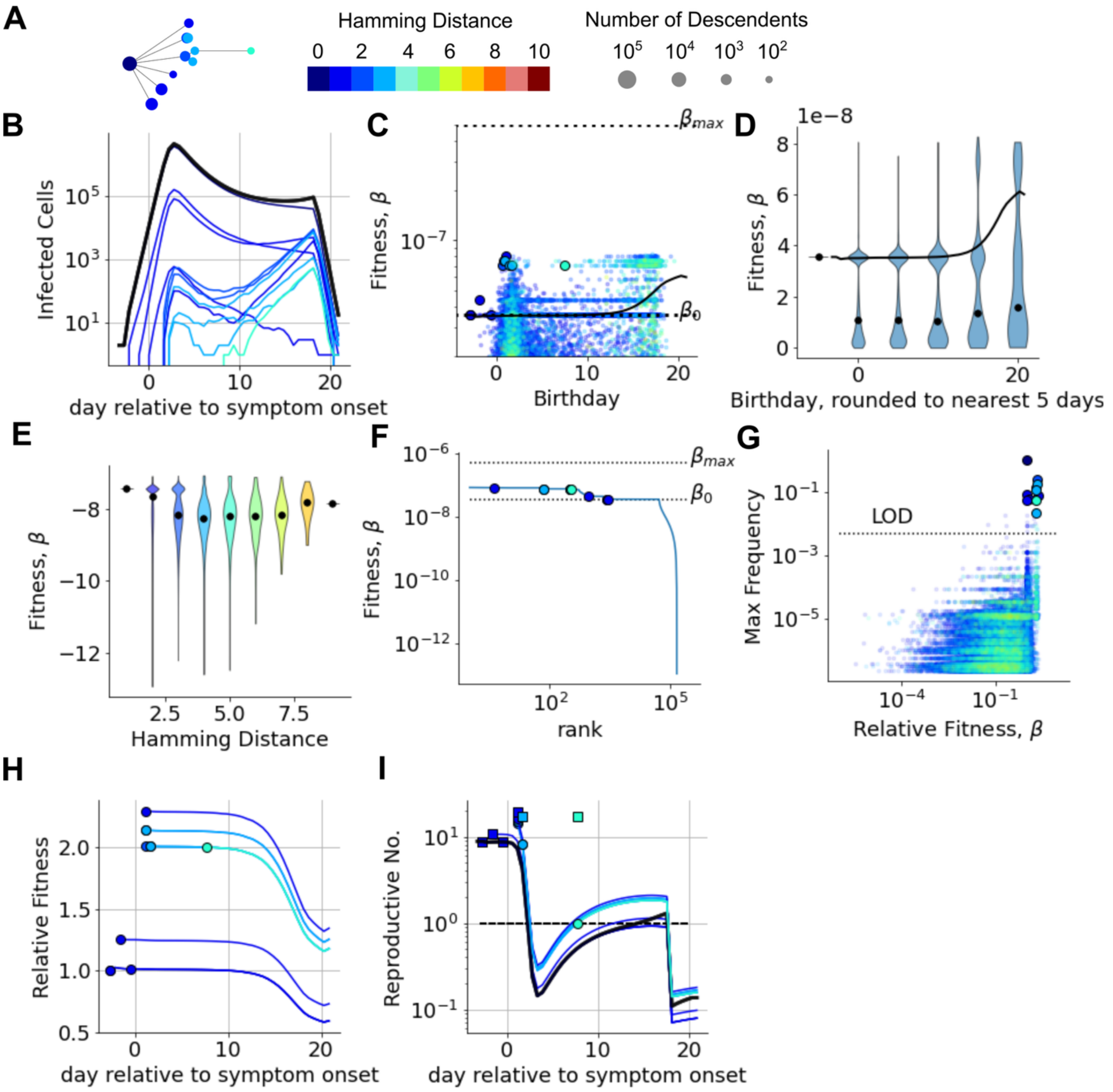


***Fig. S5: A detailed look at fitness evolution during a typical infection.*** *Panels A-I were generated using model output from one WiPhy simulation using population-calibrated evolutionary parameters and viral dynamic parameters estimated based on data from infection in one individual.*

*A) Simulated lineage of major variants (those reaching ≥2% of viral load during infection). Node radius is proportional to number of descendants (log10) and horizontal position corresponds to variant birthday (first time it appeared in simulation relative to symptom onset) concurrent with panel B. This color bar indicates HD for all panels **A-J**.*

*B) Simulated number of productive infected cells ($I_2$) corresponding to variants in **A** and their total (bold black).*

*C) Relationship between variant birthday and absolute variant fitness. Single variants (small dots colored by HD) show heterogeneity and mean (black line) trails the fitness of emerging major variants (larger dots). Dashed horizontal lines denote initial variant fitness $\beta_0$ and maximum $\beta_{max}$ fitness allowed in the simulation.*

*D) Violin plots of fitness of emerging variants grouped by 5 day intervals indicate bimodal fitness distributions with growing difference between mean (black line) and median (black dot). Note linear scale in comparison to **C**.*

*E) The bimodal fitness distributions of variants grouped by HD show increasing variance as HD increases. Median value indicated by black dot.*

*F) Each variant's fitness plotted by rank (ordered from highest to lowest infectivity). Line shows complete distribution with large dots indicating major variants. Horizontal lines following major variants reflect synonymous variants. Many variants with fitness higher than the founder never reach detectable levels.*

*G) Maximum frequency reached by variants versus their fitness relative to the population mean at time of variant appearance is roughly log log linear. Large dots indicate major variants above 2% frequency. These and other detectable minor variants (frequency above the limit of detection, LOD) have relative fitness near or above 1.*

*H) Time dynamics of the relative fitness of major variants showing that relative fitness is highest when a strain emerges and then decreases – note linear scale in comparison to* ***E*** *in which many variants have small but heterogeneous fitness.*

*I) Effective reproductive number (average number of newly infected cells from one infected cell,* ***Methods****) for each major variant (dot at birthday with line showing dynamics) compared to the population average (black line) over time after symptoms. New variants only can predominate when R>1.*

**Transmission of mutated SARS-CoV-2 variants is favored by relatively prolonged infections due to delayed immunity**

Katherine Owens, Pierce Radecki, Stefano Tempia, Anne von Gottberg, Cheryl Cohen, Eli Boritz, Joshua T. Schiffer, Daniel B. Reeves

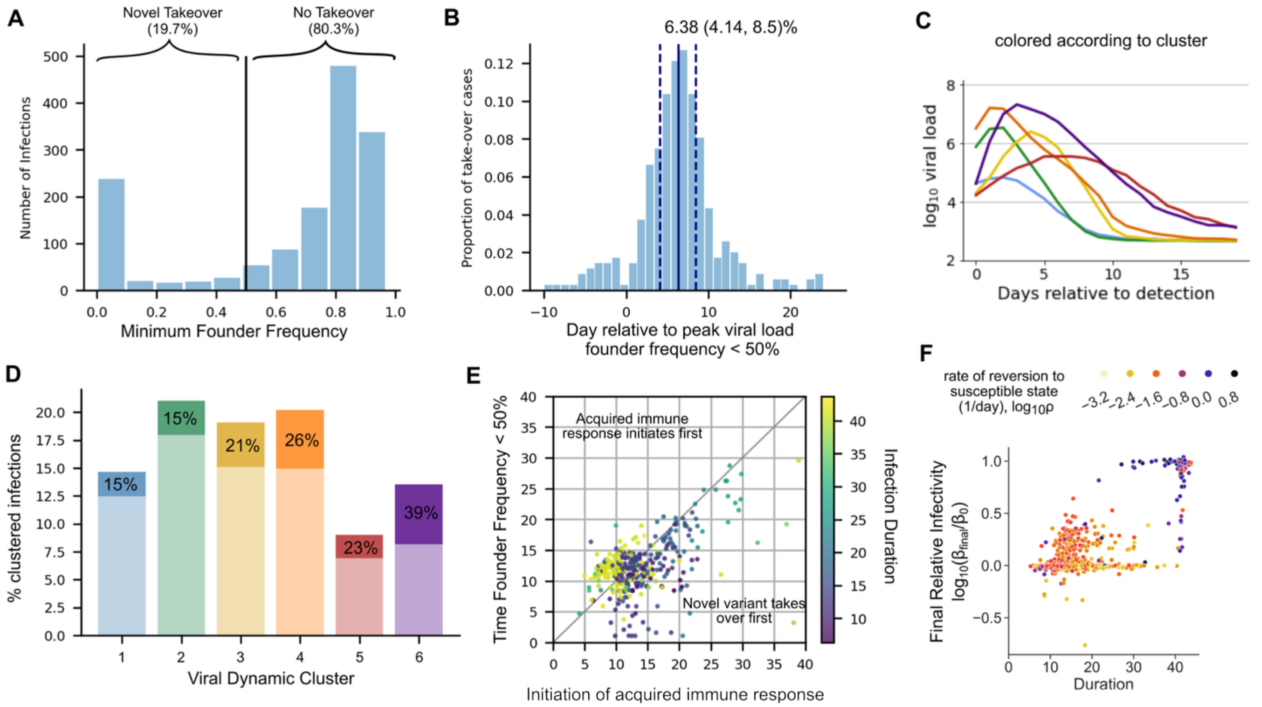


***Fig S6. Risk factors for novel variant takeover***

*A) Histogram of the minimum frequency of the founder strain observed during infection for 1452 simulations using viral dynamic parameters from individuals in the NBA cohort. Variant takeover was flagged if the founder variant's frequency dropped below 50%, which occurred in 19.7% of simulations.*

*B) Histogram of the timing of variant takeover relative to time of peak viral load for the subset of cases flagged to have takeover. This occurred a median of 6.38 days post-peak viral load (bold line) with IQR 4.14-8.5 days post-peak viral load (dashed lines).*

*C) Six characteristic viral dynamic shedding patterns identified in untreated SARS-CoV-2 infections from the NBA cohort. Reproduced from Owens et al.*

*D) Bar chart showing the percent of infections in each cluster, with light (bottom) region indicating founder-dominant simulations and saturating region (top) indicating takeover cases. The chance of a random infection from each cluster exhibiting takeover is indicated and ranges from 15% in the lowest AUC groups through 39% in the highest AUC group.*

*E) Time of takeover relative to infection vs initiation of the acquired immune response for the subset of cases flagged to have takeover. Each dot is one simulated infection and the color indicates the total duration of detectable infection.*

*F) The ratio of the average viral infectivity at the end of infection relative to the initial infectivity versus infection duration. The maximum simulation length was 40 days, so infections with a duration of 40 were considered persistent. Each dot is one simulated infection in the NBA cohort that exhibited novel variant takeover. The color indicates the the log10-transformed rate at which cells that are refractory to infection revert to a susceptible state. Infections with a high rate of reversion (purple) have a less durable innate immune response and appear more likely to have takeover lead extensive viral evolution and persistent infection. Cells with a low rate of reversion (yellow) are more likely to have a limited increase in viral fitness and clear the infection despite viral takeover occurring.*

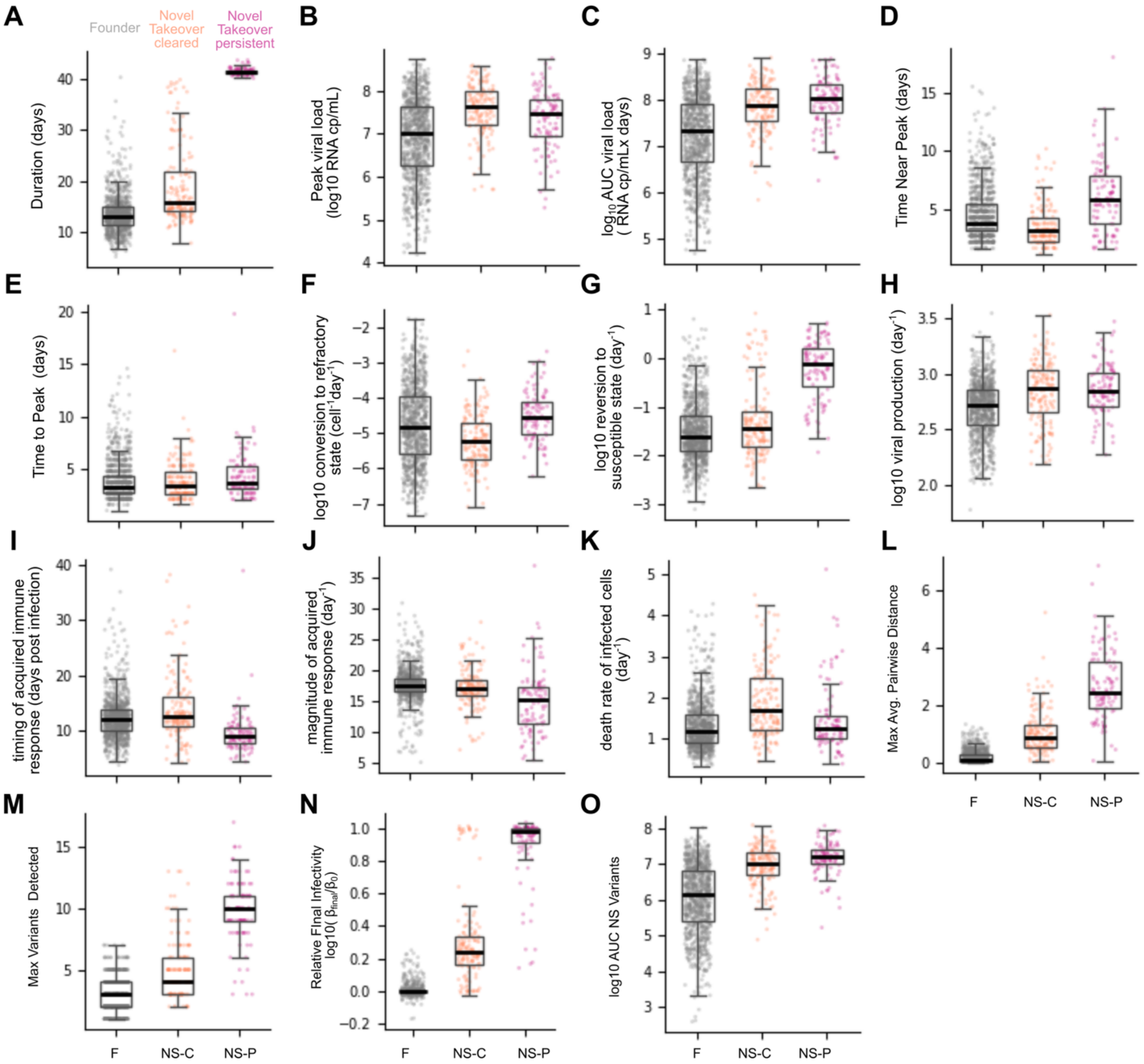


***Fig S7. Comparision of viral dynamic metrics, viral dynamic model parameter values, and viral diversity metrics between takeover groups.*** *Simulations of 1452 infections from the NBA cohort were classified as founder dominant (F) which occurred in n = 1161 cases, novel variant takeover cleared within 40 days (NS-C) which occurred in n = 191 cases, or novel variant takeover persistent for 40+ days (NS-P) which occurred in n= 100 cases. All boxplots are median, interquartile range (IQR, box) and 1.5x the IQR (whiskers).*

*A-E) Boxplots of viral dynamic metrics as defined in Table S1 including infection duration, peak viral load, log10 AUC viral load, time near peak viral load, and time to peak viral load.*

*F-K) Boxplots of viral dynamic model parameters as defined in Figure 1 including log10 rate of conversion to a refractory state ($\phi$), log10 reversion to a susceptible state ($\rho$), log10 viral production rate ($\pi$), death rate of infected cells ($\delta$), magnitude of acquired immune response ($m$), and timing of the onset of acquired immune response ($\tau$)*

*L-O) Boxplots of viral diversity metrics including maximum average pairwise distance over the course of infection, maximum number of variants detected, log-transformed ratio of infectivity of viral population at the end of infection over initial infectivity, and log10 AUC for viral load of non-synonymous variants. Founder dominated infections maintain similar viral infectivity throughout infection. Most persistent infections approach or reach the maximum permitted improvement in infectivity, 10 times the initial value.*

**Transmission of mutated SARS-CoV-2 variants is favored by relatively prolonged infections due to delayed immunity**

Katherine Owens, Pierce Radecki, Stefano Tempia, Anne von Gottberg, Cheryl Cohen, Eli Boritz, Joshua T. Schiffer, Daniel B. Reeves

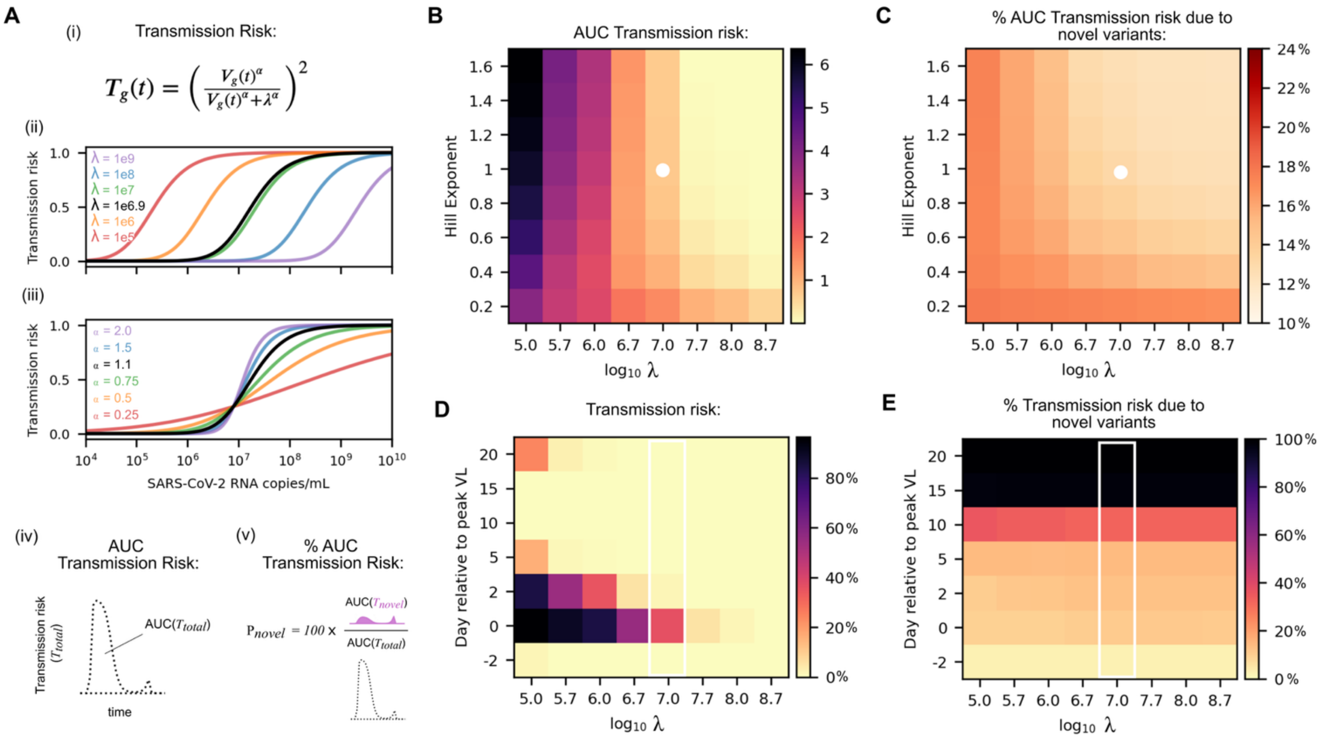


***Fig. S8: Sensitivity of projected transmission risk to risk model parameters.***

*A) We used a dose response curve from Goyal et al. and given in panel (i) to convert SARS-CoV-2 copies/mL to the risk of transmission given contact has occurred. We illustrate the dependence of this model on two parameters, $\lambda$ and $\alpha$, by plotting the predicted transmission risk for the same range of viral loads but values of $\lambda$ ranging across 4 orders of magnitude (ii) or values of alpha ranging across 2 orders of magnitude (iii). The percentage AUC transmission risk due to novel variants is calculated by applying the function from (i) to the simulated viral load at each timepoint from WiPhy model output, subsetting this total transmission risk based on the frequency of the founder strain at each time point, and dividing the AUC novel variant transmission risk curve over the total.*

*B-C) Heatmaps of AUC Transmission risk and the percentage AUC transmission risk due to novel variants for a range of hill exponents and viral loads.*

*D-E) Heatmaps total transmission risk and the percentage transmission risk due to novel variants for varying values of $\lambda$ over the course of infection. Time (vertical axis) is organized relative to peak viral load, so transmission risk is highest in every column at day 0. At this time, regardless of the choice of $\lambda$ the percentage of transmission risk due to novel variants is always around 10%.*

*G) Summary of transmission risk due to founder (84%), novel variants with increased fitness (5%), and novel variants without increased fitness (11%) across all NBA cohort infection simulations.*

***Table S1: Model Fitting Target Metrics.*** *Median and IQR are calculated among the 18 individuals from the SAI cohort used for model calibration.*

| Metric | | Definition | Median (IQR) | Units |
|---|---|---|---|---|
| *Peak viral load* | | *Maximum SARS-CoV-2 RNA measured via qPCR on nasal swab samples* | *6.8 (4.6, 8.5)* | *log10 copies/mL* |
| *Time to peak* | | *Day of first peak viral load relative to symptom onset* | *4.9 ( 0.0 , 16.0 )* | *days* |
| *Time near peak* | | *Duration viral load is continuously within 1 log of peak viral load* | *3.0 ( 1.0 , 8.0 )* | *days* |
| *Duration* | | *Day (relative to symptom onset) of first negative test following last positive test* | *7.0 (3.0, 21.0)* | *days* |
| *# variants detected* | *Day 3<br>Day 5<br>Day 10<br>Day 20* | *Number of unique variants called among SARS-CoV-2 genomes recovered* | *2 ( 1.0 , 6.0)<br>2 ( 1.0 , 6.0)<br>1.0 ( 1.0 , 6.0 )<br>1.0 (1.0,5.0)* | *count* |
| *Average Pairwise Distance* | *Day 3<br>Day 5<br>Day 10<br>Day 20* | *The total number of point mutations between each pair of sequences divided by the number of pairs in that group* | *0.02 ( 0.0 , 0.6 )<br>0.07 ( 0.0 , 0.6 )<br>0.06 ( 0.0 , 0.5)<br>0.15 ( 0.0 , 0.5 )* | *base pairs* |
| *Frequency of first dominant variant* | *Day 3<br>Day 5<br>Day 10<br>Day 20* | *The variant with the highest frequency in the first sample is the dominant variant. Frequency is the proportion of SARS-CoV-2 genomes collected that are called as that variant.* | *0.99 (0.85,1.0)<br>0.97 (0.78,1.0)<br>0.97 (0.7,0.98)<br>0.97 (0.7,0.98)* | *unitless* |

***Table S2: Viral Dynamic Parameters (reproduced from Owens et al. 2024)***

| Parameter (unit) | Symbol | Population Mean | Standard deviation of random effects | Distribution Of random effects | Source |
|---|---|---|---|---|---|
| viral infectivity ($\log_{10}$ (RNAcopies/mL)$^{-1}$ day$^{-1}$) | $\log_{10}\beta$ | -7.31 | 4.69e-2 | normal | estimated |
| viral production rate ($\log_{10}$ day$^{-1}$ ) | $\log_{10}\pi$ | 2.74 | 0.316 | normal | estimated |
| rate at which refractory cells revert to susceptible state ($\log_{10}$ day$^{-1}$ ) | $\log_{10}\rho$ | -1.62 | 0.972 | normal | estimated |
| rate constant for conversion of target cells to a refractory state ($\log_{10}$ cell$^{-1}$day$^{-1}$ ) | $\log_{10}\phi$ | -5.29 | 1.2 | normal | estimated |
| Deviation from $\log_{10}\phi$ for delta/other | $\beta_{\phi\_omicron}$ | 0.324 | -- | -- | -- |
| Deviation from $\log_{10}\phi$ for delta/other | $\beta_{\phi\_none}$ | 1.53 | -- | -- | -- |
| infected cell clearance rate when I = 1 (day$^{-1}$ cells$^{-h}$) | $\delta$ | 1.38 | 0.565 | log normal | estimated |
| onset of acquired immunity relative to detection (days) | $\tau$ | 15.9 | 0.461 | log normal | estimated |
| Deviation from $\tau$ for delta/other | $\beta_{\tau\_omicron}$ | -.364 | -- | -- | -- |
| Deviation from $\tau$ for delta/other | $\beta_{\tau\_none}$ | -.422 | -- | -- | -- |
| Deviation from $\tau$ for unvaccinated/no record | $\beta_{\tau\_>1\ vax}$ | -.141 | -- | -- | -- |
| Increase in clearance rate of infected cells due to acquired immunity (day$^{-1}$) | $m$ | 16.4 | 0.502 | log normal | estimated |
| delay between infection in nasal tissue and detection (days) | $t_0$ | 2.08 | 0.900 | logit [0,20] | estimated |
| viral clearance rate (day$^{-1}$) | $\gamma$ | 15 | -- | -- | Goyal et al. |
| mean eclipse phase duration (days$^{-1}$) | $1/k$ | 1/4 | -- | -- | Ke et al. |
